# Reduction of Nemo-like kinase increases lysosome biogenesis and ameliorates TDP-43-related neurodegeneration

**DOI:** 10.1101/2020.04.19.049395

**Authors:** Leon Tejwani, Hiroshi Kokubu, Paul J. Lee, Yangfei Xiang, Kimberly Luttik, Armand Soriano, Jennifer Yoon, Junhyun Park, Hannah Ro, Hyoungseok Ju, Clara Liao, Sofia M. Tieze, Frank Rigo, Paymaan Jafar-Nejad, Janghoo Lim

## Abstract

Protein aggregation is a hallmark of many neurodegenerative disorders, including amyotrophic lateral sclerosis (ALS). Although mutations in *TARDBP*, encoding TDP-43, account for less than 1% of all ALS cases, TDP-43-positive aggregates are present in nearly all ALS patients, including patients with sporadic ALS (sALS) or carrying other familial ALS (fALS)-causing mutations. Interestingly, TDP-43 inclusions are also present in subsets of patients with frontotemporal dementia, Alzheimer’s disease, and Parkinson’s disease; therefore, methods of activating intracellular protein quality control machinery capable of clearing toxic cytoplasmic TDP-43 species may alleviate disease-related phenotypes. Here, we identify a novel function of Nemo-like kinase (Nlk) as a negative regulator of lysosome biogenesis. Genetic or pharmacological reduction of *Nlk* increased lysosome formation and improved clearance of aggregated TDP-43. Furthermore, Nlk reduction ameliorated pathological, behavioral, and lifespan deficits in two distinct mouse models of TDP-43 proteinopathy. Because many toxic proteins can be cleared along the autophagy-lysosome axis, targeted reduction of *Nlk* represents a viable approach to therapy development for multiple neurodegenerative disorders.

## Introduction

In the majority of neurodegenerative diseases, one or more proteins aggregate over the course of disease progression and, in some cases, are thought to play a central role in pathogenesis. In the case of amyotrophic sclerosis (ALS), cytoplasmic inclusions of transactive response DNA binding protein 43 kDa (TDP-43) are observed in approximately 97% of all patients, including patients with sporadic ALS (sALS) or familial ALS (fALS) (1-3), as well as in subsets of patients with frontotemporal dementia (FTD) (1, 3), Alzheimer’s disease (4), and Parkinson’s disease (5). This aberrant accumulation of cytoplasmic TDP-43 is thought to underlie gain of toxic functions and exacerbate loss of nuclear function pathogenic mechanisms (6), therefore, preventing the formation or promoting the clearance of these inclusions may be an effective therapeutic approach for TDP-43 proteinopathies.

We have previously reported that Nemo-like kinase (Nlk) is a proline-directed serine/threonine kinase capable of interacting with and phosphorylating several neurodegenerative disease-causing proteins, namely, ataxin-1 and androgen receptor (AR), the proteins whose polyglutamine repeat expansion are causal for spinocerebellar ataxia type 1 (SCA1) and spinal and bulbar muscular atrophy (SBMA), respectively (7, 8). Genetic reduction of *Nlk* in animal models for SCA1 and SBMA alleviates several disease-related phenotypes, and this partial rescue was previously attributed to decreased phosphorylation of two Nlk substrates, ataxin-1 and AR, at key residues involved in the pathogenic mechanisms (7, 8). Although it is known that Nlk interacts with several other proteins that are expressed within the central nervous system, the physiological role of Nlk and the general effects of its modulation in the adult nervous system have not yet been comprehensively examined.

In this study, we identify a novel function of Nlk as a negative regulator of lysosome gene transcription in the nervous system. Reduction of Nlk promoted functional lysosome biogenesis through increasing the expression of a large number of genes involved in major aspects of lysosomal function, while its overexpression suppressed this transcription. Overexpression of Nlk promoted the accumulation and aggregation of TDP-43 and its cleavage products that are normally cleared through the autophagy-lysosome pathway (ALP), while reduction of Nlk enhanced clearance of aggregated TDP-43 *in vitro* and *in vivo* in a transgenic mouse model of TDP-43- related fALS. Furthermore, genetic or pharmacological reduction of *Nlk* using antisense oligonucleotides (ASOs) alleviated motor behavioral and pathological deficits in two distinct TDP- 43 mouse models. The data presented here uncover and characterize a novel approach to enhancing lysosomal clearance of aggregated proteins relevant to ALS/FTD and can potentially be applied to a broad range of neurodegenerative diseases.

## Results

### Nlk negatively regulates the lysosome

Beyond its role in altering phosphorylation levels of ataxin-1 and AR, the disease-causing proteins for SCA1 and SBMA, respectively, we sought to interrogate the cellular functions of Nlk in the nervous system. To this end, we utilized dual guide RNAs to target Cas9 nickase to *Nlk* in Neuro2a (N2a) cells, generating two offset single-stranded nicks for error-prone non-homologous end-joining repair (Figure 1A) (9). Due to the polyploidy of N2a cells, we assessed the efficacy of mutagenesis by examining residual Nlk protein levels (Supplemental Figure 1A). Individual clones were classified by amount of intact Nlk protein, and to ensure that the phenotypes observed were a result of Nlk depletion and not spurious artifacts of the CRISPR/Cas9 mutagenesis, multiple distinct clones with undetectable Nlk levels (*Nlk* KO) were expanded and used for further experimentation (Supplemental Figure 1B). RNA sequencing (RNA-seq) was performed on mRNA from triplicates of *Nlk* KO cells and wild-type (WT) isogenic controls to identify transcriptional changes caused by *Nlk* reduction in an unbiased fashion (Supplemental Figure 2A,B, and Supplemental Table 1). Interestingly, cellular component enrichment analyses on all differentially expressed genes revealed an enrichment for transcriptional changes related to multiple cytoplasmic membrane-bound vesicles, including endosomes and lysosomes (Figure 1B), but not to stress granules that are membraneless structures implicated in ALS pathogenesis (10) (Supplemental Figure 2C,D). Additionally, lysosomal storage disorders were among the significant disease pathways highlighted by the enrichment analysis (Figure 1C).

**Figure 1.**
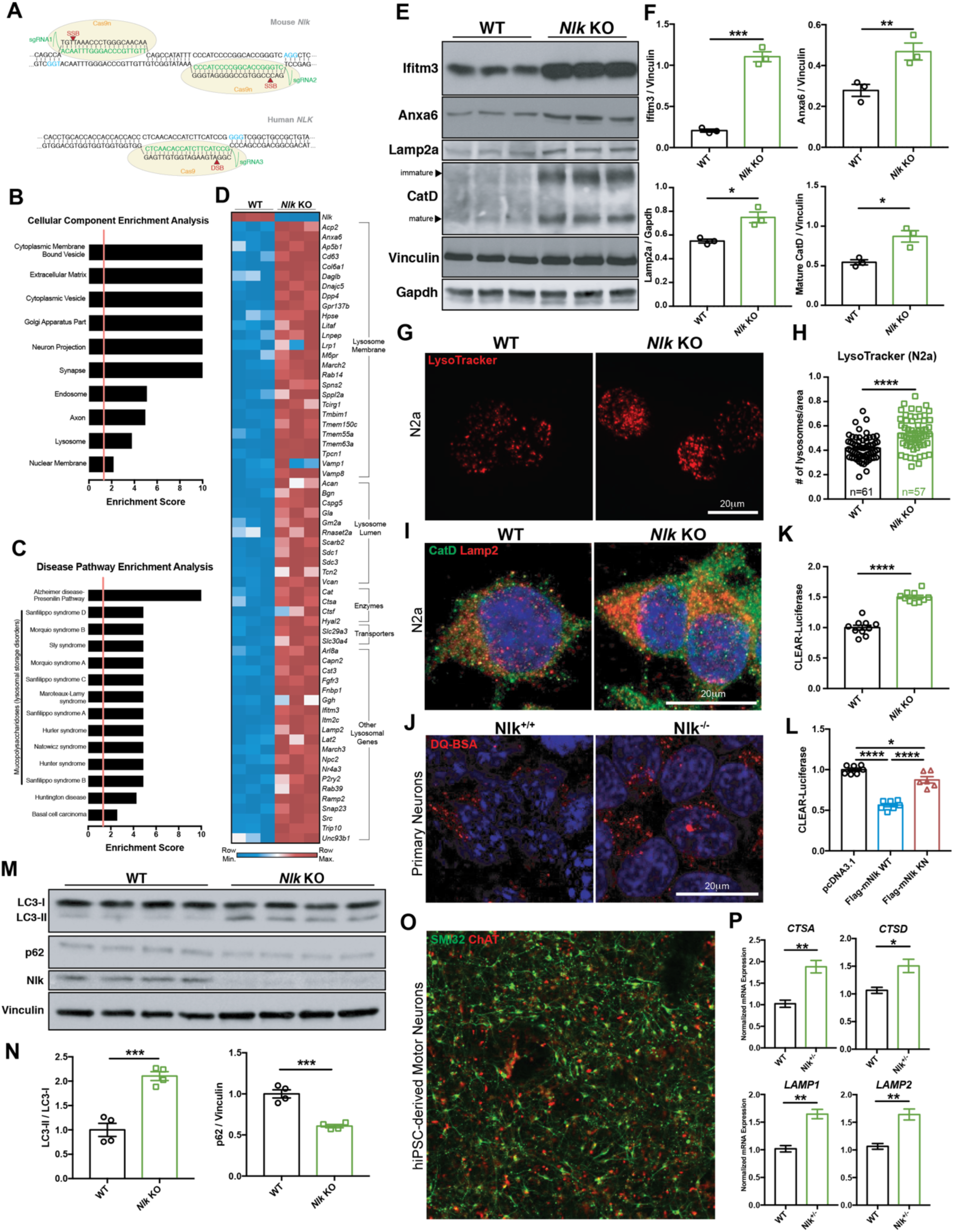
Reduction of *Nlk* increases lysosome biogenesis *in vitro*. (**A**) Schematic of dual guide RNA targeting of Cas9 nickase (Cas9n) to *Nlk* for generation of isogenic *Nlk* knock-out (KO) N2a cells and *NLK*^*+/-*^ human iPSCs. (**B**,**C**) Enrichment analyses for cellular components (**B**) and disease pathways (**C**) for all significantly differentially expressed genes from RNA-seq analysis of isogenic WT and *Nlk* KO N2a cells. Significant enrichment scores of 1.3 (FDR adjusted p-value q<0.05) are designated by vertical red lines. (**D**) Heatmap of significantly up-regulated lysosomal genes from RNA-seq. (**E**,**F**) Western blot validation (**E**) confirmed RNA-seq results, quantified in (**F**). (**G**,**H**) Representative Lysotracker images of WT and *Nlk* KO N2a cells (**G**), quantified in (**H**) demonstrated increased lysosome number in *Nlk* KO cells. (**I**) Immunofluorescence imaging of Lamp2^+^ and CatD^+^ lysosomes in WT and *Nlk* KO N2a cells. (**J**) Immunofluorescence imaging of functional lysosomes in DIV7 primary cortical neurons incubated with DQ-BSA. (**K**,**L**) CLEAR-luciferase assay in *Nlk* KO N2a cells (**K**) or WT N2a cells transfected with WT or kinase-negative (KN) Nlk (**L**) demonstrated Nlk kinase activity suppresses CLEAR network transcription. (**M**,**N**) Western blots (**M**) showed increased LC3-II/LC3-I ratio and decreased p62 levels in *Nlk* KO N2a cells, quantified in (**N**). (**O**) Representative immunostaining images of human iPSC-derived motor neurons (**o**). (**P**) qPCR data showing *NLK*^*+/-*^ iPSC-derived motor neurons expressed higher levels of lysosomal genes *CTSA, CTSD, LAMP1*, and *LAMP2* compared to isogenic controls. Two-tailed t-tests or one-way ANOVA analyses were performed and mean±s.e.m are displayed. *p<0.05, **p<0.01, ***p<0.001, ****p<0.0001.

**Figure 2.**
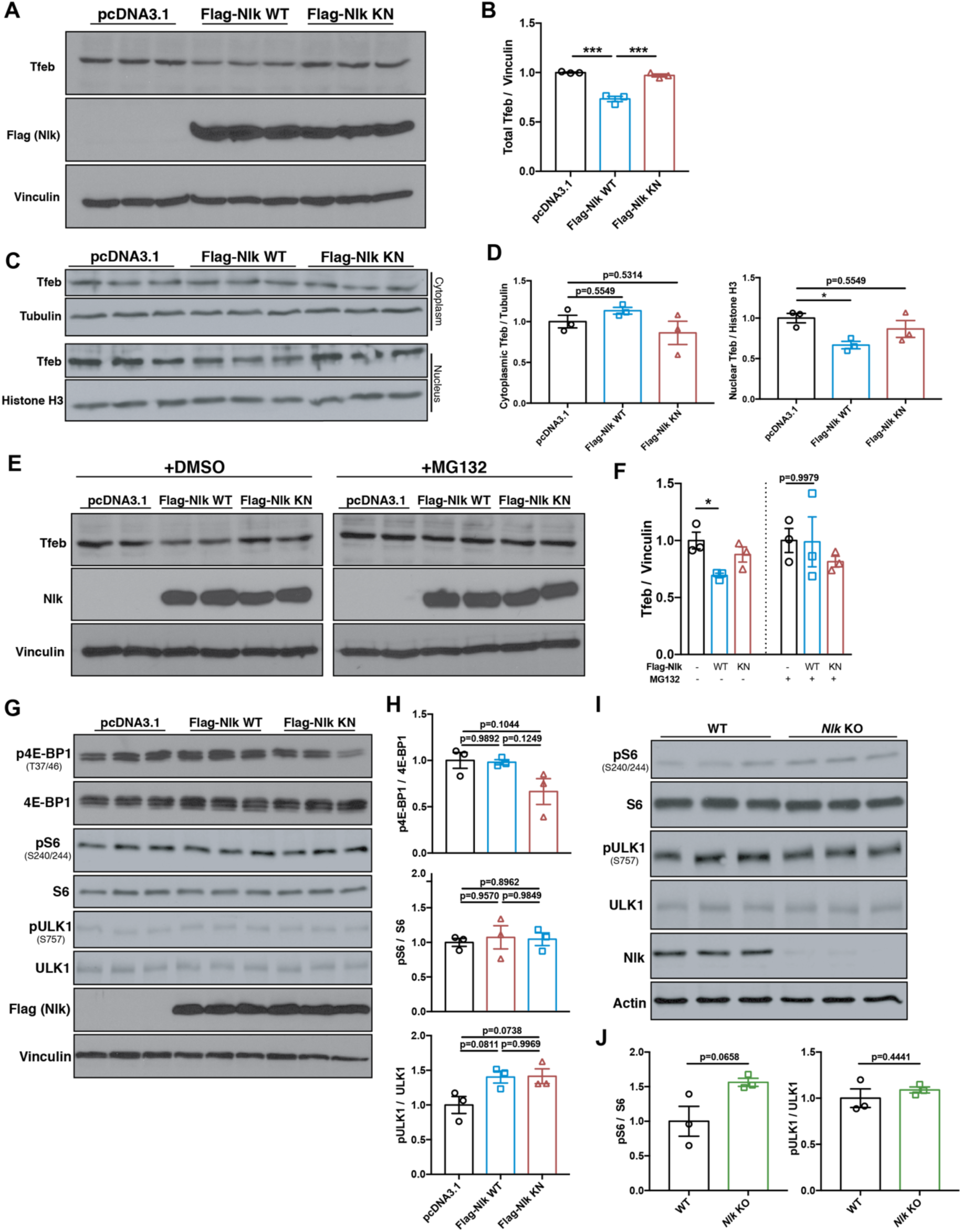
Nlk de-stabilizes nuclear Tfeb via the proteasome without affecting mTOR activity. (**A**,**B**) Western blots of N2a whole-cell protein lysates following transfection with Flag-Nlk WT or Flag- Nlk KN (**A**). Tfeb levels decreased upon Nlk overexpression in a kinase activity-dependent manner, quantified in (**B**). (**C**,**D**) Western blots of WT N2a cells transfected with Flag-Nlk WT or Flag-Nlk KN showed a kinase activity-dependent decrease of nuclear Tfeb levels (**C**), quantified in (**D**). (**E**,**F**) Western blots of N2a whole-cell protein lysates following transfection with Flag-Nlk WT or Flag- Nlk KN, with DMSO vehicle or MG132 treatment (**E**). Tfeb levels decreased upon Nlk overexpression in a kinase activity-dependent manner, but this destabilization was not observed in the presence of the proteasome inhibitor MG132, quantified in (**F**). (**G**,**H**) Western blots of N2a whole-cell protein lysates showed transfection with Flag-Nlk WT or Flag-Nlk KN did not affect p4E-BP1/4E-BP1, pS6/S6, or pULK1/ULK1 ratios, suggesting mTOR independence. (**I**,**J**) Western blots of N2a whole-cell protein lysates showed *Nlk* KO did not affect pS6/S6 or pULK1/ULK1 ratios. Two-tailed t-tests or one-way ANOVA analyses were performed and mean±s.e.m are displayed. *p<0.05, ***p<0.001.

Because of the overrepresentation of up-regulated lysosome-associated transcripts in the *Nlk* KO dataset (Figure 1D), some of which we validated at the protein level (Figure 1E,F), we sought to identify the functional consequences of Nlk reduction on lysosome biogenesis and function. We determined that lysosome number was increased in *Nlk* KO N2a cells compared to isogenic controls by performing LysoTracker live imaging and immunostaining for lysosomal markers Lamp2 and Cathepsin D (Figure 1G-I). Transfection of exogenous Nlk into Nlk-deficient N2a cells suppressed this increase in lysosome number in a kinase activity-dependent manner (Supplemental Figure 3A). Addition of DQ-BSA, a self-quenched dye that becomes fluorescent upon proteolysis by lysosomal hydrolases, to primary cortical neurons from *Nlk*^*+/+*^ and *Nlk*^*-/-*^ mice revealed increased fluorescence in the absence of Nlk (Figure 1J), suggesting that not only does the number of lysosomes increase, but their proteolytic functionality does as well. Furthermore, transfection of a 4XCLEAR-Luciferase reporter (11) confirmed that *Nlk* KO promoted transcriptional activity of genes belonging to the Coordinated Lysosomal Enhancement and Regulation (CLEAR) network of autophagy and lysosomal genes (12, 13) in multiple distinct clones (Figure 1K and Supplemental Figure 3B), while overexpression of Nlk suppressed 4XCLEAR-Luciferase reporter activity in a kinase activity-dependent manner (Figure 1L). Using a tandem fluorescent-tagged RFPGFP-LC3 reporter (14) (Supplemental Figure 3C), we observed that *Nlk* KO increased the delivery of reporter-containing autophagosomes to the lysosome, which was inhibited by overexpression of WT Nlk (Supplemental Figure 3D). LC3-II to LC3-I ratio was increased with a concomitant decrease in p62 in *Nlk* KO cells (Figure 1M,N), while overexpression of WT Nlk led to an accumulation of p62 (Supplemental Figure 3E). Finally, human motor neurons differentiated from CRISPR/Cas9-mutated *NLK*^*+/-*^ induced pluripotent stem cells (iPSCs) (Figure 1A and Supplemental Figure 4) expressed higher levels of lysosomal genes *CTSA, CTSD, LAMP1*, and *LAMP2* compared to isogenic WT control cells (Figure 1O,P). Collectively, these data demonstrate that *Nlk* reduction increases autophagic flux and cargo degradation by increasing lysosome biogenesis, while *Nlk* overexpression impairs lysosomal degradation of autophagy substrates.

**Figure 3.**
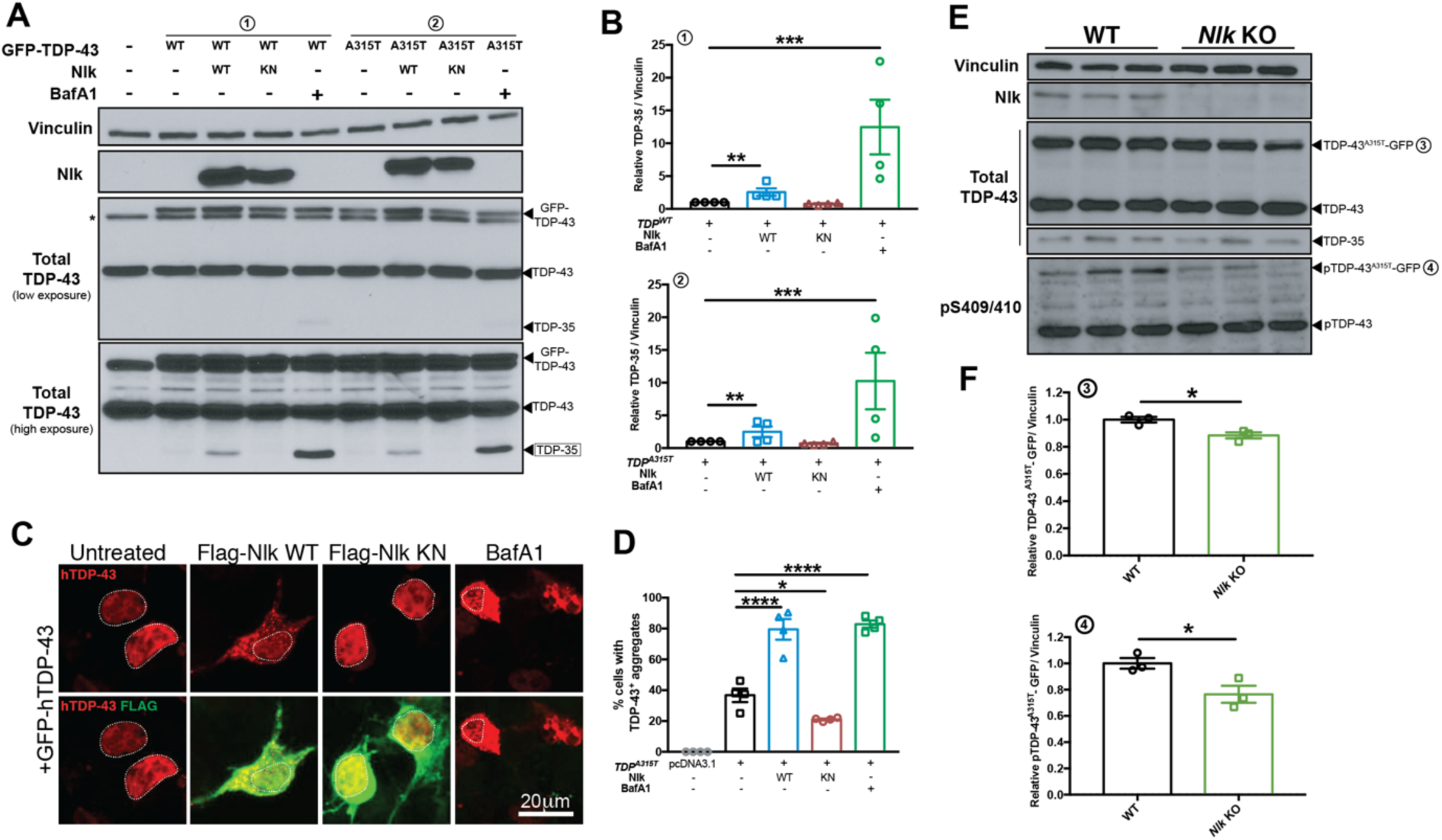
Genetic reduction of *Nlk* reduces TDP levels *in vitro*. (**A**,**B**) Western blots showing co-expression of Nlk-WT with GFP-tagged TDP^WT^ or TDP^A315T^ in NSC-34 cells (**A**) increased levels of TDP-35, while Nlk-KN did not. V-ATPase inhibitor bafilomycin A1 (BafA1) treatment similarly increased TDP-35 levels in the absence of Nlk overexpression. Quantification of TDP- 35 levels in (**B**). (**C**,**D**) Representative immunostaining of NSC-34 cells co-transfected with TDP-43 and Nlk displaying Nlk-WT increased formation of cytoplasmic TDP aggregates (**C**), quantified in (**D**). Nuclei are outlined with dotted lines. (**E**,**F**) Western blots of cell lysates from WT or *Nlk* KO N2a cells transfected with TDP-43^A315T^-GFP (**E**), quantified in (**F**). Levels of exogenous TDP-43-GFP and pTDP-43-GFP were significantly reduced. Two-tailed t-tests were performed between the two genotypes. *p<0.05, **p<0.01, ***p<0.001, ****p<0.0001. One-way ANOVA analyses were performed to compare all listed genotypes/treatments unless otherwise noted and mean±s.e.m are displayed.

**Figure 4.**
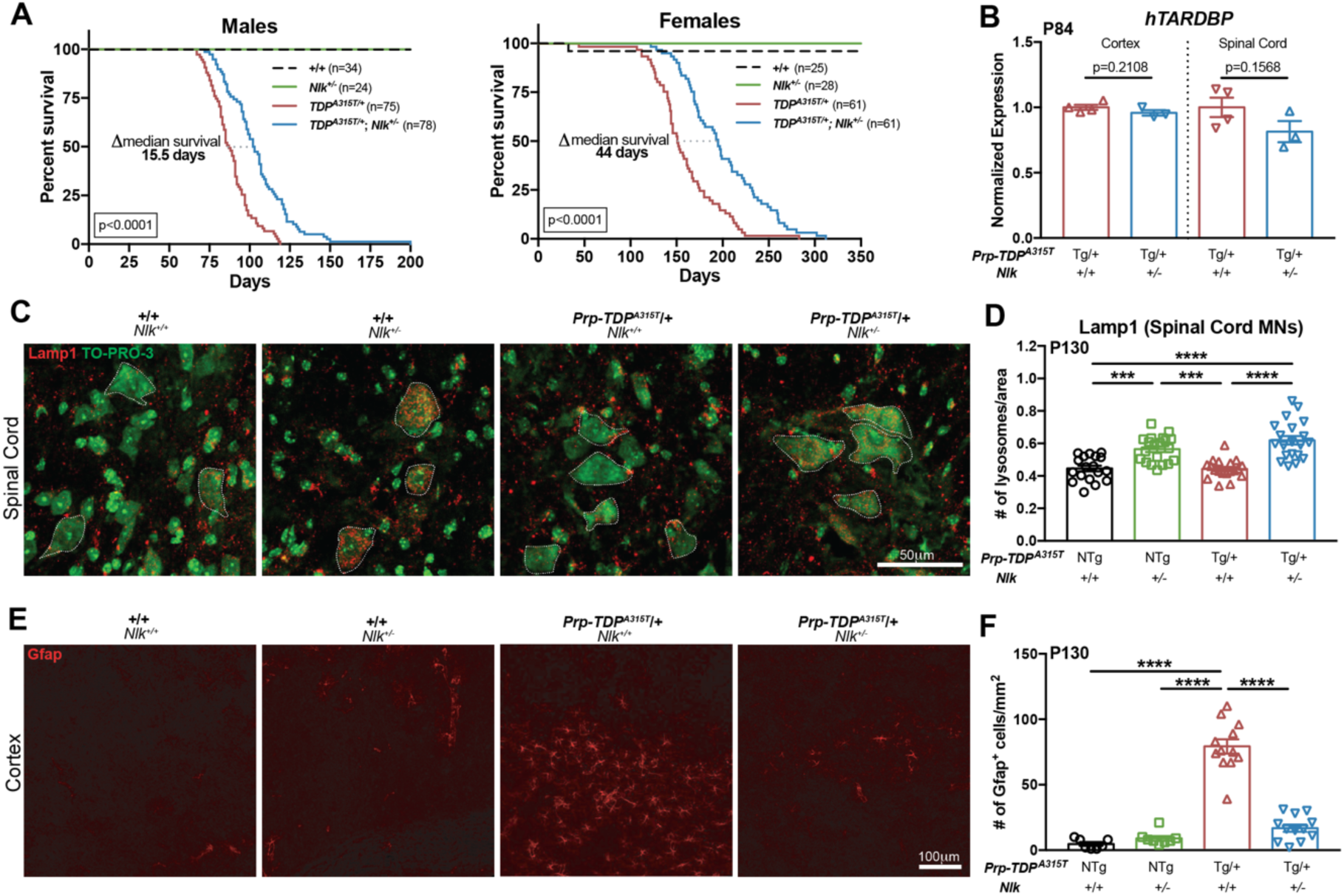
Genetic reduction of *Nlk* improves survival and pathology in TDP-43 mice. (**A**) Kaplan-Meier survival curves showing 50% genetic reduction of *Nlk* increased *Prp-TDP*^*A315T/+*^ male and female animal survival. Data are combined from two independent *Nlk* gene trap insertion lines. Curves were compared by log-rank test. (**B**) 50% genetic reduction of *Nlk* did not affect cortex or spinal cord *hTARDBP* mRNA level in P84 *Prp- TDP*^*A315T/+*^ mice. (**C-F**) Representative immunohistochemical images showing genetic reduction of *Nlk* increased Lamp1^+^ lysosomes in the soma of lumbar spinal cord motor neurons (**C**, quantified in **D**; n=motor neurons) and rescued layer V astrogliosis (**E**, quantified in **F**; n=sections) in P130 *Prp-TDP*^*A315T/+*^ female mice. Two-tailed t-tests were performed between the two genotypes. One-way ANOVA analyses were performed to compare all listed genotypes/treatments unless otherwise noted and mean±s.e.m are displayed. NTg= non-transgenic, Tg= transgenic. *p<0.05, **p<0.01, ***p<0.001, ****p<0.0001.

### Nlk destabilizes nuclear Tfeb

Several transcription factors have been reported to be involved in the transcriptional regulation of a large number of lysosomal genes and among these transcription factors, Tfeb has been described as a key master regulator of lysosome gene expression and function (12, 13, 15). To determine if Nlk exerts its control of lysosome-associated genes through altering Tfeb, we assessed levels of Tfeb in N2a cells in which WT or kinase-negative (KN) Nlk was overexpressed. In whole cell lysates, total Tfeb levels were reduced with Nlk overexpression in a kinase activity-dependent manner (Figure 2A,B). Because the ability of Tfeb to induce lysosomal gene expression is highly dependent on its translocation to the nucleus (16), we next isolated nuclear and cytoplasmic fractions to assess changes in the subcellular localization of Tfeb. Interestingly, overexpression of Nlk induced a nucleus-specific reduction in Tfeb levels, with no changes in cytoplasmic levels (Figure 2C,D), suggesting the nuclear depletion of Tfeb was not attributed to its cytosolic retention. Furthermore, because the reduction in Tfeb was compartment-specific (Figure 2C,D) and *Tfeb* transcript levels were not significantly altered in Nlk-deficient N2a cells (Supplemental Table 1), this regulation was not a result of transcriptional changes of *Tfeb* itself.

Because the nuclear reduction of Tfeb appeared to post-translational and not due to a shuttling deficiency from the cytoplasm, we next examined the contribution of the ubiquitin-proteasome system (UPS), a major protein catabolism pathway active in both the cytoplasm and nucleus involved in the turnover of short- and long-lived proteins (17). Treatment with the proteasome inhibitor MG132 normalized total Tfeb levels following Nlk overexpression (Figure 2E,F). Therefore, we conclude that the Nlk-induced Tfeb destabilization in the nucleus is dependent on the proteasome. Phosphorylation of Tfeb at specific residues by various kinases, the best documented being mTOR, has been ascribed to be a major mechanism that governs subcellular localization and activity of Tfeb (16, 18-21). To test if this novel mechanism of Tfeb regulation by Nlk could similarly be mediated through a direct effect of Nlk on mTOR signaling, we examined several key molecules in the mTOR signaling cascade. We determined that *Nlk* KO or overexpression of Nlk in N2a cells did not affect 4E-BP1, S6, or ULK1 phosphorylation (Figure 2G-J), suggesting the observed effect on Tfeb by Nlk is likely not through altering overall mTOR activity. Collectively, these data reveal a novel mechanism for the regulation of Tfeb levels by Nlk, and a new component of the intricate cross-talk between the transcriptional control of the lysosome and the UPS that has not yet been previously described.

### Nlk regulates TDP-43 levels

The ALP is one of two primary systems for intracellular protein degradation and has been implicated to be defective in some forms of ALS (22, 23). In most neurodegenerative disorders, including ALS, one or more proteins aberrantly accumulate, and these aggregates are typically associated with disease pathogenesis. Although the genetic basis underlying ALS is heterogeneous (2), ubiquitin-positive TDP-43 inclusions are present in nearly all ALS patients (1, 3). Because soluble and aggregated forms of TDP-43 and its C-terminal fragment (CTF) cleavage products can be cleared through the ALP (24, 25), we aimed to determine if TDP-43 levels could be modulated through altering Nlk levels. Co-expression of TDP-43^WT^ or mutant TDP-43^A315T^, a fALS-causing variant of TDP-43 (26), with Nlk in motor neuron-like NSC-34 cells induced a kinase activity-dependent increase in TDP-43-positive inclusions and cleavage product TDP-35, both of which also increased with lysosome inhibitor Bafilomycin A1 (BafA1) (Figure 3A-D). Conversely, *Nlk* reduction in N2a cells reduced protein levels of exogenous TDP-43 and pS409/410 TDP-43 (Figure 3E,F), a disease-specific form present across TDP-43 proteinopathies (27). This reduction in pS409/410 TDP-43 was likely not a result of a decrease in direct phosphorylation by Nlk, as Nlk did not physically interact with TDP-43 under the tested conditions (Supplemental Figure 5). Furthermore, Nlk is a proline-directed serine/threonine MAPK-like kinase not capable of phosphorylating at S409 or S410. Collectively, these data suggest an enhanced indirect clearance mechanism of pathological TDP-43 by Nlk.

**Figure 5.**
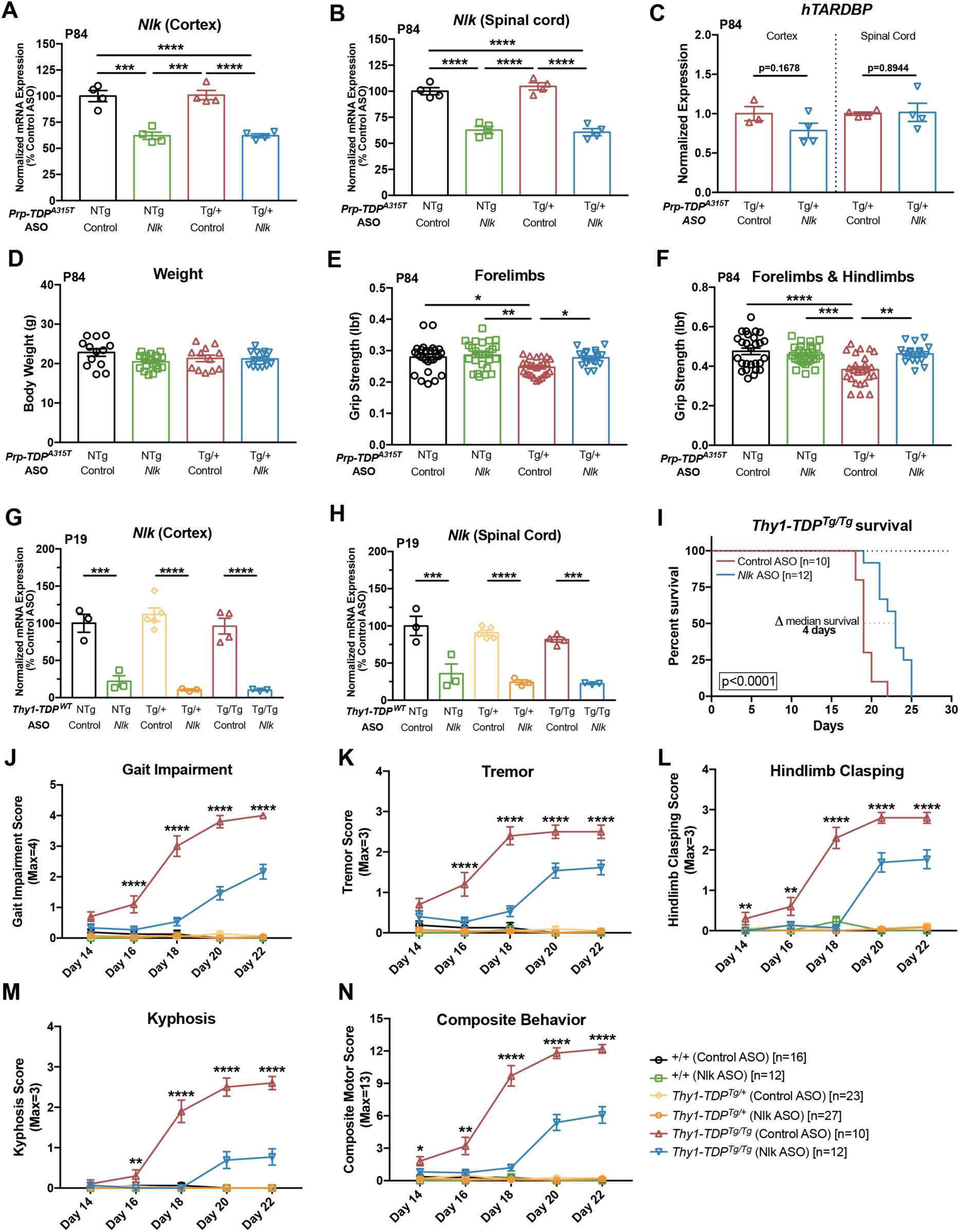
Pharmacological reduction of *Nlk* using ASOs improves behavior in two TDP-43 mouse models. (**A**,**B**) ICV injection of 10μg of *Nlk* ASO at P1 resulted in a 40% reduction of *Nlk* mRNA at P84 in the cortex (**A**) and spinal cord (**B**). (**C**) ASO reduction of *Nlk* did not affect transcript levels of *hTARDBP*. (**D**) Overall body weight was not affected by ASO administration. (**E**,**F**) *Nlk* ASO administration at P1 rescued forelimb (**E**) and forelimb with hindlimb (**F**) grip strength deficits in *Prp-TDP*^*A315T/+*^ mice at P84. (**G**,**H**) Injection of 10μg of *Nlk* ASO at P1 resulted in a 90% and 75% reduction of *Nlk* mRNA at P19 in the cortex (**G**) and spinal cord (**H**), respectively. (**I**) Kaplan-Meier survival curves showing administration of 10μg *Nlk* ASO increased *Thy1-TDP*^*Tg/Tg*^ animal survival. Curves were compared by log-rank test. (**J-N**) *Nlk* ASO administration reduced gait impairment (**J**), tremor (**K**), hindlimb clasping (**L**), kyphosis (**M**), and composite motor score (**N**) in *Thy1-TDP*^*Tg/Tg*^ animals between P14-P22. One-way ANOVA analyses were performed to compare all listed genotypes/treatments per day unless otherwise noted and mean±s.e.m are displayed. *p<0.05, **p<0.01, ***p<0.001, ****p<0.0001.

### Genetic reduction of *Nlk* extends TDP-43 mouse survival

We hypothesized that increasing lysosome biogenesis and TDP clearance through targeted Nlk reduction could modify pathogenesis in animal models of TDP-43 proteinopathies. To test this, we crossed two independent *Nlk* gene trap loss-of-function mouse lines (*Nlk*^*XN619/+*^ and *Nlk*^*RRJ297/+*^) (7, 8), collectively referred to here as *Nlk*^*+/-*^, with transgenic mice expressing a fALS-causing mutant *hTARDBP*^*A315T*^ under the control of a *Prp* promoter (28), referred to here as *Prp-TDP*^*A315T/+*^. Although male and female *Prp-TDP*^*A315T/+*^ hemizygous mice display differences in survival that are likely due to sex differences in gastrointestinal complications that are associated with lethality (29), genetic reduction of *Nlk* by 50% increased median survival of male and female *Prp- TDP*^*A315T/+*^ mice by about 20% (Figure 4A and Supplemental Figure 6A,B), without altering *hTARDBP* transcript levels (Figure 4B). In addition to increasing overall survival, disease onset, which was defined as the first time point in which weight gain was no longer observed (30, 31), was significantly delayed in male mice with a 50% genetic reduction of *Nlk* (Supplemental Figure 6C). Examination of postnatal day (P)130 female mice revealed that genetic reduction of *Nlk* was able to increase Lamp1-positive lysosome number in motor neurons of the lumbar spinal cord *in vivo* (Figure 4C,D). Finally, astrogliosis in layer V of the motor cortex was reduced to near-control levels in *Prp-TDP*^*A315T/+*^ mice with *Nlk* reduction (Figure 4E,F). These data demonstrate that *Nlk* reduction *in vivo* causes an increase in lysosome biogenesis that, presumably through increasing clearance of toxic TDP-43 species, alleviates pathological changes and extends lifespan of *Prp- TDP*^*A315T/+*^ hemizygous mice.

**Figure 6.**
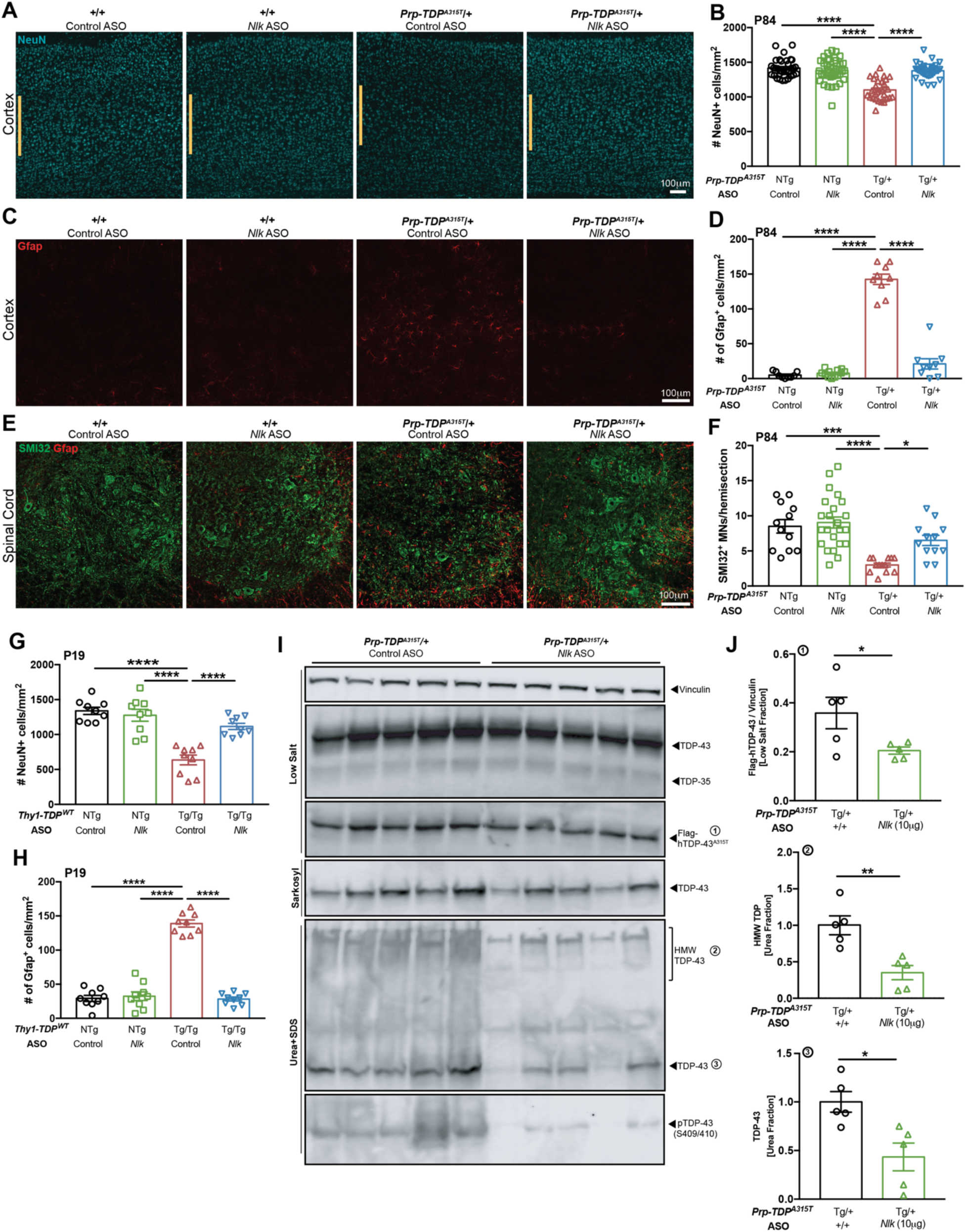
Pharmacological reduction of *Nlk* using ASOs decreases TDP-43 levels *in vivo* and ameliorates pathology in two TDP-43 mouse models. (**A-B**) Administration of 10μg of *Nlk* ASO at P1 rescued loss of layer V cortical neurons (**A**, quantified in **B**; n=sections), layer V astrogliosis (**C**, quantified in **D**; n=sections), and lumbar spinal cord motor neuron loss (**E**, quantified in **F**; n=sections) in *Prp-TDP*^*A315T/+*^ male mice at P84. (**G**,**H**) 10μg of *Nlk* ASO at P1 rescued loss of layer V cortical neurons (**G**; n=sections) and layer V astrogliosis (**H**; n=sections) in *Thy1-TDP*^*Tg/Tg*^ mice at P19. (**I**,**J**) Western blots of various forms of TDP-43 in fractions of male mouse spinal cords at P84. Quantification of TDP species from (**I**) shows levels of Flag-hTDP-43 (low salt fraction), total TDP-43 (urea fraction), and high-molecular weight TDP (urea fraction) were significantly reduced in *Nlk* ASO injected animals (**J**). Two-tailed t-tests were performed between the two genotypes. One-way ANOVA analyses were performed to compare all listed genotypes/conditions unless otherwise noted and mean±s.e.m are displayed. *p<0.05, ***p<0.001, ****p<0.0001.

### *Nlk* ASOs improve TDP-43 mouse behavior

To validate the approach of *Nlk* reduction for neurodegenerative diseases in a more translationally-relevant manner, we generated ASOs targeting the mouse *Nlk* transcript. ASOs are short oligonucleotides that hybridize to complementary transcripts with high specificity and target them for RNase H-mediated degradation (32). The chemically-modified backbone and bases minimize immunogenicity and enhance ASO stability, enabling persistent knockdown of a target transcript for prolonged periods of time. In recent years, ASO-based approaches have shown promise in pre- clinical models of various motor neuron diseases (33-35), as well as in human clinical trials for spinal muscular atrophy (36) and Huntington’s disease (37). Intracerebroventricular (ICV) injections of *Nlk* ASOs at P1 led to a dose-dependent reduction of *Nlk* mRNA levels in the cortex and spinal cord (Figure 5A,B and Supplemental Figure 7A,B) with no overt astrogliosis or microgliosis at P84 (Supplemental Figure 7C-F). Administration of *Nlk* ASOs to *Prp-TDP*^*A315T/+*^ mice did not affect levels of *hTARDBP* transcript (Figure 5C) or overall body weight (Figure 5D).

Based on our dosing experiments (Supplemental Figure 7A,B), we selected a dose of 10μg *Nlk* ASO, which produced a robust and persistent knockdown efficiency of *Nlk* by 40% at P84 following a single P1 injection (Figure 5A,B). We began by assessing the effect of *Nlk* ASOs on motor function at P84, a time point at which *Prp-TDP*^*A315T/+*^ mice display brain and spinal cord pathology (28). *Prp-TDP*^*A315T/+*^ mice displayed reduced forelimb and forelimb with hindlimb grip strength compared to non-transgenic controls, and these deficits were restored to control levels in transgenic animals that received *Nlk* ASOs (Figure 5E,F).

To confirm the benefit of *Nlk* reduction on motor deficits associated with TDP-43 accumulation, we turned to an additional transgenic mouse model that overexpresses WT *hTARDBP* under the control of a *Thy1* promoter (38), referred to here as *Thy1-TDP*^*Tg/Tg*^. Compared to the *Prp-TDP*^*A315T/+*^ mice, these animals display much more severe behavioral deficits characterized by a rapidly progressing motor impairment and eventual paralysis (necessitating euthanasia at a humane endpoint) when bred to give rise to homozygous progeny (38). P1 ICV delivery of 10μg of *Nlk* ASO reduced *Nlk* mRNA levels at P19 by 90% and 75% in the cortex and spinal cord, respectively (Figure 5G,H). Median lifespan in *Thy1-TDP*^*Tg/Tg*^ animals injected with *Nlk* ASOs at P1 was increased by 21% compared to control ASO injected animals (Figure 5I). Additionally, motor behavior was scored as previously described (34, 39) and *Nlk* reduction ameliorated gait impairment, tremor, hindlimb clasping, and kyphosis between P14-P22 (Figure 5J-N, Supplemental Figure 8, and Supplemental Video 1). Finally, because altering diet may affect the course of disease in transgenic ALS mouse models (40), we raised an independent cohort of *Thy1-TDP*^*Tg/Tg*^ animals on a high fat/gel food diet and observed a similar improvement in motor behavior with a greater extension of survival with P1 *Nlk* ASO injections (Supplemental Figure 9).

### *Nlk* ASOs reduce TDP-43-related pathology

We next examined the effect of pharmacological reduction of *Nlk* on pathological and biochemical changes in TDP-43 transgenic mice. *Prp-TDP*^*A315T/+*^ mice displayed a loss of about 22% of layer V neurons of the motor cortex at P84 with an accompanying astrogliosis, both of which could be normalized to near control levels with a single P1 injection of 10μg *Nlk* ASO (Figure 6A-D, Supplemental Figure 10A,B). Similarly, loss of lumbar spinal cord SMI32-positive ventral horn motor neurons was mitigated with *Nlk* reduction (Figure 6E,F, Supplemental Figure 10C). The pathological rescue with *Nlk* ASOs was also observed in P19 *Thy1-TDP*^*Tg/Tg*^ animals, as layer V cortical neuron loss and astrogliosis were prevented (Figure 6G,H, and Supplemental Figure 10D,E).

Our *in vitro* experiments describing the effect of Nlk reduction to promote lysosomal function and clearance of TDP-43 (Figures 1,3), along with the confirmation that genetic reduction of *Nlk* increases lysosome gene expression and number in mouse and human iPSC-derived spinal cord motor neurons (Figure 1P, and Figure 4C,D), suggest that the therapeutic benefit we observe *in vivo* could be mediated through altering levels of aggregated TDP-43 in motor neurons. To confirm that *Nlk* reduction did indeed alter levels of aggregated TDP-43 *in vivo*, we sequentially isolated protein fractions based on detergent solubility (3) from spinal cords of male P84 *Prp- TDP*^*A315T/+*^ mice that had been injected with control or *Nlk* ASOs at P1. Biochemical analysis of TDP-43 confirmed that *Nlk* reduction using ASOs decreased overall TDP-43 levels in the more insoluble fractions, with the most striking differences observed in the highly insoluble urea fraction (Figure 6I,J). Importantly, *Nlk* reduction did not affect soluble wild-type TDP-43 levels (Figure 6I), suggesting a preservation of TDP-43 to engage in its normal, critical functions. Collectively, these data demonstrate that *Nlk* reduction can confer beneficial therapeutic effects in multiple murine models of protein aggregation.

## Discussion

The inability to adequately clear aggregation-prone proteins is a common phenomenon across neurodegenerative disorders. In ALS, a vast majority of patients display toxic cytoplasmic TDP- 43-positive inclusions, regardless of sporadic or familial etiology. Therapeutic approaches aimed at preventing inclusion formation or enhancing clearance are viable strategies; however, methods that globally reduce TDP-43 levels are unfavorable, as TDP-43 serves essential functions in the nucleus (6). On the other hand, increasing activity of cellular machinery capable of clearing cytoplasmic species of aggregated TDP-43, such as the ALP, may be a more effective approach, as this may reduce toxic effects in the cytoplasm while also potentially restoring the proper distribution of TDP-43 to the nucleus (41).

Here, we have identified Nlk as a novel negative regulator of the lysosome, whose modulation can affect transcription of lysosome-associated genes, and therefore, alter the capacity to clear disease-associated aggregated proteins, such as TDP-43, through the ALP without affecting levels of soluble wild-type TDP-43 species. To test the therapeutic potential of this approach, we employed genetic and pharmacological tools to specifically reduce *Nlk* in two mouse models of TDP-43 proteinopathy and observed substantial lifespan, behavioral, biochemical, and pathological rescue. Furthermore, we have identified a novel molecular mechanism for the regulation of Tfeb through the nuclear proteasome. It is possible that Nlk reduction also affects additional target molecules in parallel with Tfeb, such as other transcription factors that may directly alter lysosome gene expression. This study warrants further investigation into these downstream and parallel molecules, which may reveal novel targets of interest that confer greater precision when designing therapeutic agents.

We have previously reported that NLK phosphorylates AR and ataxin-1, the proteins mutated to cause SBMA and SCA1, respectively, and that phosphorylation of these proteins contributes to pathogenicity (7, 8). Rather than a direct effect of altering phosphorylation of the disease-causing protein, the mechanism of therapeutic benefit by Nlk reduction described here is an indirect one likely mediated through increased lysosomal clearance of aggregated TDP-43, as there is no evidence for a physical interaction between NLK and TDP-43. It is possible that this role of NLK as a negative regulator of lysosome biogenesis described here may, in part, contribute to the amelioration of disease phenotypes upon genetic reduction of *Nlk* in SCA1 and SBMA animals, beyond simply altering phosphorylation of the toxic protein. A further dissection of the effect of these direct and indirect pathways on suppressing mutant ataxin-1 and AR toxicity could provide critical insights into the role of Nlk in neurodegenerative diseases and whether there is a unifying protective effect of Nlk reduction.

The therapeutic benefits of Nlk reduction on motor behavior, neurodegeneration and its associated astrogliosis, and survival of two independent TDP-43 animal models provide strong proof-of-principle evidence for this promising approach in TDP-43 proteinopathies, both in instances in which *TARDBP* mutations are present (a subset of ALS patients), and when *TARDBP* mutations are not present but TDP-43 aggregation occurs. However, because genetic and pharmacological reduction of *Nlk* occurred either prenatally or at P1 in our animal models, respectively, the observed improvements may be attributed to alterations in the disease onset and/or progression. Our data demonstrate that constitutive genetic reduction of Nlk by 50% delays the median onset of disease by 10.5 days in male *Prp-TDP*^*A315T/+*^ mice. Similarly, pharmacological reduction using *Nlk* ASOs delays the onset of motor deficits in *Thy1-TDP*^*Tg/Tg*^ animals. Because the median survival of male *Prp-TDP*^*A315T/+*^; *Nlk*^*+/-*^ animals increases by 15.5 days compared to transgenic animals with wild-type Nlk levels, it is possible that Nlk reduction not only delays the onset, but also slows progression of disease. These findings warrant a more thorough future examination of pathological alterations comparing animals at the onset, early disease, and late stages of disease to definitively determine if disease progression can also be altered by Nlk reduction.

Due to the precision afforded by complementary binding of an ASO to its target transcript of interest, ASOs have emerged as an invaluable tool in specifically altering levels and/or processing of a variety of proteins involved in neurodegenerative diseases, most of which have been the disease-causing protein itself (33, 42-45). While these approaches have proven to be extremely effective in preclinical genetic models of these disorders, the utility of such approaches is restricted to small subsets of patients that carry specific, rare mutations in the gene of interest. In the case of ALS, the vast majority of patients have a sporadic form of the disease in which a genetic etiology is not identifiable; therefore, targeted reduction of a mutated protein is not possible. For this reason, alternative approaches that mitigate pathologies commonly observed across sALS and fALS, such as pathological TDP-43 aggregation (34) or premature poly-adenylation and loss of Stathmin-2, a key regulator of microtubule dynamics in motor neurons (46, 47), are highly desirable.

To our knowledge, the present study is one of the first to generate and validate the therapeutic benefit of ASOs targeting a disease modifier (and not the disease-causing gene itself) in rodent models of a neurodegenerative disease. Among these, we believe this study is the first to target a novel molecule that has not been implicated as a genetic risk factor for the disease being studied. Because no known neurological condition in humans has been associated with a reduction of Nlk, mice with partial genetic reduction of *Nlk* display no pathological (neurodegenerative or inflammatory), survival, or cognitive changes up to 52 weeks (latest time point analyzed; data not shown), and mice with postnatal whole-body conditional ablation of *Nlk* are grossly normal (data not shown), we believe targeted reduction of *Nlk* or its downstream effectors to increase ALP function is a potentially generalizable therapeutic strategy for several neurodegenerative disorders, beyond just ALS. Further examination of the appropriate and effective time window for therapeutic intervention in specific disease contexts would be necessary prior to proceeding.

## Materials and Methods

### Animal husbandry

All animal procedures were performed in accordance with the National Institutes of Health Guide for Care and Use of Experimental animals and approved by the Yale University Institutional Animal Care and Use Committee. Mice were maintained on a 12 hours light and 12 hours dark cycle with standard mouse chow, or high fat breeder chow followed by DietGel food (ClearH_2_O), and water *ad libitum*. Two independent *Nlk* gene trap mouse lines (*Nlk*^*RRJ297/+*^ and *Nlk*^*XN619/+*^) (7), referred to simply as *Nlk*^*+/-*^, were maintained on a pure C57BL/6J background, as previously described. We refer to compound heterozygous (*Nlk*^RRJ297/XN619^) mice as *Nlk* knockout (*Nlk*^-/-^). Transgenic B6.Cg-Tg(Prnp-TARDBP*A315T)95Balo/J (*Prp-TDP*^*A315T/+*^) mice were obtained from The Jackson Laboratory (Stock #010700) and maintained on a pure C57BL/6J background. Transgenic B6;SJL-Tg(Thy1-TARDBP)4Singh/J (*Thy1-TDP*^*Tg/+*^) hemizygous mice were obtained from The Jackson Laboratory (Stock #012836) and maintained on a mixed B6;SJL background. For animal PCR genotyping, tail snips for DNA extraction were taken for *Thy1-TDP*^*Tg*^ animals at P14, and between P18-P20 for all other animals. Both male and female mice were used for behavior, survival, and pathological analyses. Unless otherwise noted, both male and female mice were analyzed.

### Primary neuronal cultures

For primary cortical neuron culture, mouse cortex was dissected from postnatal day 0.5 (P0.5) mouse brains in ice-cold HBSS and washed three times with HBSS. Tissue was dissociated in Papain (Worthington) for 20 minutes at 4°C. After dissociation, tissue was washed with HBSS three times and further dissociated with P200 and P1000 pipette tips in Neurobasal medium (Gibco) supplemented with 10% B27 (Gibco), 2mM GlutaMAX-I (Gibco), 1mM Sodium Pyruvate (Gibco), and 1% Penicillin/Streptomycin (Gibco). Cortical neurons were plated in 6-well plates on coverslips coated with 100μg ml^-1^ poly-D-lysine (Sigma-Aldrich) and incubated at 37°C with 5% CO_2_. On the next day, full culture medium was changed and subsequently, half media was changed every three days. Seven days *in vitro* (DIV7) neurons were used for DQ-BSA experiments.

### N2a and NSC-34 cell lines

N2a and NSC-34 cells were cultured in Dulbecco’s modified Eagle’s medium (DMEM) (Gibco), supplemented with 10% (v/v) fetal bovine serum (FBS) (Gibco). Transient transfection was performed using Lipofectamine 2000 (ThermoFisher) according to manufacturer’s instruction. Cells were analyzed 48 hours after transfection, unless otherwise noted. Cells were routinely tested for mycoplasma contamination by PCR as previously described (48) using primers For-GGCGAATGGGTGAGTAACACG and Rev- CGGATAACGCTTGCGACCTATG. Any mycoplasma-positive samples were immediately discarded. For RFPGFP-LC3 experiments, ptfLC3 (Addgene #21074) (14) was transfected using Lipofectamine 2000, as described above.

### N2a *Nlk* CRISPR cell line generation

For generation of N2a *Nlk* CRISPR cell lines, CRISPR guide RNA sequences (sgRNA#1 TTGTTGCCCAGGGTTTAACA and sgRNA#2 CCCATCCCCGGCACCGGGTC) targeting the mouse *Nlk* locus were cloned into pSpCas9n(BB)-2A-Puro (Addgene #62987; PX462). Cells were transfected with gRNA and Cas9n-expressing plasmids using Lipofectamine 2000 and successfully transfected cells were selected with 3μg ml^-1^ puromycin for two days. Single clones were generated by plating cells at a single cell density in 96-well plates and then expanded to 24-well plates for genotyping by western blotting.

### hiPSC *NLK* CRISPR cell line generation

For generation of hiPSC *NLK* CRISPR cell lines, CRISPR guide RNA sequence (sgRNA#3 CTCAACACCATCTTCATCCG) targeting the human *NLK* locus was cloned into pSpCas9 (BB)-2A-GFP (Addgene #481387 PX458). Wild-type iPSCs were transfected with gRNA and Cas9-expressing plasmid using an Amaxa Nucleofector 2b and successfully transfected cells were collected by FACS, as previously described(48). Single clones were generated by plating cells at a single cell density and then expanded to 24-well plates for genotyping by Sanger sequencing and western blotting.

### Human iPSC-derived motor neuron differentiation

Human motor neurons were generated using a previously-established protocol with minor modifications (49). Briefly, hiPSC colonies were split with Dispase (1U/ml) (Stem Cell Technologies) and re-plated at 1:6 ratio on Matrigel-coated plate in mTeSR1 medium. The next day (day 1), mTeSR1 medium was replaced with neural medium supplemented with 3μM CHIR99021 (Tocris), 2μM DMH1 (Tocris), and 2μM SB431542 (Stemgent). The neural medium contained DMEM/F12 and Neurobasal medium mixed at 1:1 ratio, 0.5 × N2, 0.5 × B27, 100μM ascorbic acid (Sigma), 1 × Glutamax (Life Technologies), 1 × penicillin/streptomycin (Life Technologies). The medium was replenished on day 3 and day 5. On day 7, neuroepithelial progenitors (NEPs) were split with Dispase (1U/ml) and re-plated at 1:6 ratio on Matrigel-coated plate in neural medium supplemented with 1μM CHIR99021 (Tocris), 2μM DMH1 (Tocris), 2μM SB431542 (Stemgent), 0.1μM retinoic acid (RA) (Stemgent), 0.5μM Purmorphamine (Stemgent). The medium was replenished on day 9, 11, and 13. On day 14, motor neuron progenitors (MEPs) were dissociated with Accutase (Stem Cell Technologies) and re-plated at 1:6 ratio on Matrigel-coated plate in neural medium supplemented with 3μM CHIR99021 (Tocris), 2μM DMH1 (Tocris), 2μM SB431542 (Stemgent), 0.1μM retinoic acid (RA) (Stemgent), 0.5μM Purmorphamine (Stemgent), and 0.5 mM VPA (Stemgent). The medium was replenished on day 16, 18 and 20. On day 21, MEPs were dissociated with Accutase and re-plated at 1:6 ratio on Matrigel-coated plate in neural medium supplemented with 0.5μM retinoic acid (RA) (Stemgent) and 0.1μM Purmorphamine (Stemgent). The medium was replenished on day 23 and 25. From day 27, Compound E (Calbiochem) was supplemented to the above medium to facilitate motor neuron maturation, and media changes were performed every other day. Mature motor neurons were dissociated with Accutase and plated on Matrigel-coated 35-mm glass bottom dish or Matrigel-coated coverslip for immunostaining.

### RNA extraction, sequencing, and analysis

RNA was extracted using the Qiagen RNeasy Mini Kit and genomic DNA was removed with DNase I as recommended in the manufacturer’s instructions. The total RNA was sent to the Yale Center for Genome Analysis for processing. RNA integrity was measured using an Agilent Bioanalyzer 2100 (Agilent Technologies) and RIN values were assessed to ensure high RNA integrity before sequencing. Libraries were generated using oligo-dT purification of polyadenylated RNA, followed by reverse transcription into cDNA prior to being fragmented, blunt ended, and ligated to adaptors. The library was quantified before pooling and sequencing on an Illumina HiSeq 2000 using a 75 bp paired-end read strategy. All sequencing FASTQ files and processed normalized output can be found under GEO accession # xxxxx.

TopHat2 v2.1.0 was utilized to align reads to the mouse reference genome (GRCm38/mm10) before quantification and differential expression analysis with Cufflinks v2.2.1 (50-53). Cuffnorm was utilized for generating normalized expression values (52) and annotations with a False Discovery Rate (FDR) adjusted p-value (q<0.05) were considered significant. Enrichment analysis was carried out using Gene Set Enrichment Analysis (GSEA) (54, 55), ToppCluster (56), and Ingenuity Pathway Analysis (IPA, Qiagen Winter Release 2019) on all differentially regulated genes. Biological pathways and cellular components with enrichment scores greater than 1.3 (-log Bonferroni-corrected p-value for ToppCluster, and -log Benjamini-Hochberg corrected p-value for IPA) were considered significant. The diseases function of Toppcluster, which links different gene expression profiles to those known for specific disorders, was utilized to investigate any diseases that have an over-represented number of genes differentially regulated in the *Nlk* KO cells. The Gene Ontology Consortium (GO) (57, 58) was utilized to generate gene lists for lysosome, autophagy, and stress granule-related genes. Enrichment score bar plots and heatmaps were plotted in R version 3.3.3.

For real-time quantitative PCR (qPCR), cDNA was synthesized using oligo-dT primers and iScript cDNA synthesis kit (Biorad). qPCRs were run using Taqman probes with iTaq Universal Probe Supermix on a C1000 Thermal Cycler (BioRad) equipped with BioRad CFX Manager software. The following Taqman probes (Applied Biosystems) were used: *LAMP1* (Hs00931461_m1), *LAMP2* (Hs00174474_m1), *CTSA* (Hs00264902_m1), *CTSD* (Hs00157205_m1), *HPRT1* (Hs02800695_m1), *Nlk* (Mm00476435_m1), *TARDBP* (Hs00606522_m1), *Gfap* (Mm01253033_m1), *Aif1* (Mm00479862_g1), *Hprt* (Mm03024075_m1), and *Actb* (#4352933E). Expression data were determined by normalizing target expression to housekeeping genes (*Hprt* and *Actb*) using BioRad CFX Manager software and then plotted using Prism 7 (GraphPad).

### Protein extraction and western blot analysis

Western blot was performed as described previously (8). For protein extraction from cells, cells were harvested and lysed in Triple lysis buffer (0.5% NP-40, 0.5% Triton X-100, 0.1% SDS, 20mM Tris-HCl (pH 8.0), 180mM NaCl, 1mM EDTA, and Roche cOmplete protease inhibitor cocktail) for 15 minutes at 4°C, rotating and then centrifuged for 10 minutes at 13,000rpm at 4°C.

For Nlk co-immunoprecipitation experiments, cell pellets were harvested in NP-40 lysis buffer (0.5% NP-40, 20mM Tris (pH 8.0), 180mM NaCl, 1mM EDTA, and Roche cOmplete protease inhibitor cocktail and PhosSTOP protease inhibitors) for 15 minutes at 4°C, rotating and then centrifuged for 10 minutes at 13,000rpm at 4°C. A fraction of the resulting supernatant was set aside to be used as the input fraction. The remaining supernatant was incubated with anti-FLAG M2 magnetic beads overnight at 4°C. On the following day, the bound fraction was collected and analyzed by SDS-PAGE.

Protein extraction from mouse tissue for TDP-43 analysis was performed as previously described(3), with minor modifications. Briefly, frozen tissue was weighed and dounce homogenized in 5mL/g LS buffer (10mM Tris pH 7.5, 5mM EDTA, 1mM DTT, 10% sucrose, cOmplete EDTA-free protease inhibitors and PhosSTOP protease inhibitors) (Roche). Samples were ultracentrifuged at 25,000xg for 30 minutes at 4°C and the supernatant was collected as the “Low Salt” fraction. The pellet was washed again with LS buffer, ultracentrifuged at 25,000xg for 30 minutes at 4°C, the supernatant was discarded, and the pellet was then resuspended in HS buffer (LS buffer, 1% Triton X-100, 500mM NaCl). The supernatant of this fraction was collected as the “High Salt” fraction. The pellet was resuspended in MF buffer (HS buffer, 30% sucrose) to remove myelin. Following ultracentrifugation at 180,000xg for 30 minutes at 4°C, the pellet was resuspended in SK buffer (LS buffer, 1% sarkosyl, 500mM NaCl) and solubilized at 22°C for 60 minutes using a ThermoMixer 5350 (Eppendorf) at 500rpm. The solubilized mixture was then ultracentrifuged at 180,000xg for 30 minutes at 22°C. The supernatant was collected as the “Sarkosyl” fraction. The remaining pellet was resuspended in urea/SDS buffer (30mM Tris-HCl, 7M urea, 2M thiourea, 2% SDS). This fraction was ultracentrifuged at 25,000xg for 30 minutes at 22°C and the supernatant was isolated as the “Urea” fraction.

Total protein concentrations of isolated protein lysates were quantified by BCA protein assay kit (ThermoFisher) and equal protein amounts were boiled at 95°C for 10 minutes, loaded, and analyzed by SDS-PAGE. Protein from gels were transferred onto PVDF or nitrocellulose membranes overnight at 4°C. The next day, membranes were washed three times, blocked by 5% non-fat dry milk in TBST for one hour at room temperature, followed by incubation with primary antibody in 1% non-fat dry milk in TBST at 4°C overnight. Membranes were washed with TBST three times and incubated with sheep anti-mouse or donkey anti-rabbit IgG conjugated with horseradish peroxidase (HRP) (1:4,000, GE Healthcare) for 1 hour at room temperature. Membranes were developed using Western Lightning Plus-ECL reagent (PerkinElmer) and visualized using a SRX-101A tabletop X-ray film processor (Konica Minolta) or KwikQuant Imager (Kindle Biosciences).

The following primary antibodies were used: rabbit anti-IFITM3 (ProteinTech; 11714-1-AP; 1:2,000), rabbit anti-Annexin VI (Abcam; ab31026; 1:1,000), mouse anti-Vinculin (Sigma-Aldrich; V9264; 1:10,000), rabbit anti-Lamp2a (Abcam; ab18528; 1:1,000), rabbit anti-CatD (Abcam; ab75852; 1:2,000), mouse anti-Gapdh (Sigma-Aldrich; G8795; 1:10,000), rabbit anti-LC3B (Abcam; ab51520; 1:3,000), mouse anti-p62 (Novus; H00008878-M01; 1:10,000), rabbit anti-NLK (Abcam; ab26050; 1:1,000), rabbit anti-TFEB (MyBioSource; MBS004492; 1:500), mouse anti-Tubulin (Sigma-Aldrich; T6557; 1:10,000), rabbit anti-Histone H3 (Millipore; #05-928; 1:5,000), rabbit anti-TARDBP (Novus; NB110-55376; 1:1,000), rabbit anti-phospho-TDP-43 Ser409/410-2 (Cosmo Bio Co; TIP-PTD-M01; 1:1,000), mouse anti-Flag (Sigma-Aldrich; F3165; 1:10,000), rabbit anti-HA (abcam; ab9110; 1:5000); rabbit anti-4E-BP1 (Cell Signaling; #9644, 1:1,000), rabbit anti-phospho-4E-BP1 (Cell Signaling; #2855; 1:1,000), rabbit anti- ULK1 (Cell Signaling; #8054; 1:1000), rabbit anti-phospho-ULK1 (Cell Signaling; #14202; 1:1000), rabbit anti-S6 (Cell Signaling; #2217; 1:1000), and rabbit anti-phospho-S6 (Cell Signaling; #2215; 1:1000).

### Cytoplasmic and nuclear fractionation

N2a cells transfected with indicated plasmids were lysed in fractionation buffer containing 0.5% Triton X- 100, 50mM Tris-HCl, 137.5mM NaCl, 10% Glycerol, and 5mM EDTA and incubated on ice for 20 minutes, followed by centrifugation for three minutes at 1,000xg at 4°C. The supernatant was then transferred to a new Eppendorf tube and centrifuged again for 5 minutes at 15,000rpm at 4°C. This supernatant was used as a cytoplasmic fraction. The pellet from the first centrifugation was washed with fractionation buffer twice and then suspended in 0.5% SDS in 100mM Tris-HCl. These samples were sonicated and centrifuged for 10 minutes at 13,000rpm at 4°C. The supernatant after this centrifugation was used as a nuclear fraction. Cytoplasmic and nuclear fractions were quantified by BCA protein assay kit and analyzed by SDS-PAGE and western blotting, as described above.

### Animal tissue collection and cryo-sectioning

Mice were euthanized using isoflurane and perfused with ice-cold PBS for three minutes to remove blood. Spinal cords were then extracted by hydraulic extrusion, as described previously (59). Briefly, mice were decapitated and skin along the mouse back was removed. The spinal column was cut transversely at the pelvic bone. A syringe loaded with cold PBS was inserted into the caudal spinal column and the spinal cord was extruded using hydraulic force. The thoracic and sacral portions of the spinal cord were immediately flash frozen using dry ice with isopropanol and used for protein and RNA extraction, respectively. Fresh lumbar cord tissue was post-fixed in cold 4% paraformaldehyde (PFA) for 48 hours, and cryopreserved by immersion in a 20% to 30% sucrose gradient, followed by freezing in OCT compound (VWR). Fresh brains were dissected and cut into sagittal halves. The right brain was post-fixed for 48 hours and cryo-preserved using a 20% to 30% sucrose gradient, followed by freezing in OCT compound. The left brain was macro-dissected and individual tissues were flash frozen using dry ice with isopropanol, then stored at -80°C until further processing.

Embedded and frozen lumbar spinal cords were serially sectioned transversely as 30μm sections directly onto SuperFrost Plus slides (ThermoFisher) using a Cryostat CM1850 (Leica). Slides with spinal cord sections were stored at -80°C until further use. Slides with spinal cord sections from comparable segments were identified using gross anatomy under a light microscope and selected for further analysis. Embedded and frozen brains were sectioned using a Cryostat CM1850 as 30μm free-floating sagittal sections in PBS. Floating sections were stored in PBS at 4°C until immunostaining was performed.

### Immunostaining and microscopy

Cultured cell lines and primary cortical neurons were fixed with 4% PFA in PBS for 10 minutes at room temperature followed by washing three times with PBS and incubation with 5% normal goat serum (NGS) for one hour at room temperature. Immunocytochemical staining was performed by over-night incubation with primary antibodies at 4°C. On the next day, primary antibody was removed and cover slips were washed with PBS three times and incubated with Alexa488-, Alexa555-, Alexa568-, Alexa594-, or Alexa633-conjugated secondary antibodies (1:500, Invitrogen) for two hours at room temperature, protected from light. Cells were counterstained with TO-PRO-3 (1:5,000, Invitrogen) together with secondary antibodies when necessary. Immunostained cells were mounted onto slides using Vectashield mounting reagent (Vector Laboratories). For immunohistological staining of mouse tissue, sections were washed three times with 0.25% Triton X-100 (American Bioanalytical) in PBS (PBS-X), blocked with 5% NGS in PBS-X for 1 hour, followed by over-night incubation with primary antibodies at 4°C. On the next day, sections were washed three times with PBS-X, incubated with conjugated secondary antibodies (1:500, Invitrogen) for two hours at room temperature, protected from light. Sections were then washed three times with PBS-X, once with PBS, and mounted using Vectashield mounting reagent with DAPI (Vector Laboratories).

For lysotracker imaging, N2a cells were incubated with 100nM Lysotracker Red DND-99 (Invitrogen) in a 5% CO_2_ incubator at 37°C for 55 minutes followed by three washes with HBSS. Live imaging was then performed using an UltraVIEW VoX (Perkin Elmer) inverted spinning disc confocal microscope equipped with a 60x CFI Plan APO VC, NA 1.4, oil objective and Volocity acquisition software (Improvision). Cells were maintained in Live Cell Imaging Solution (Invitrogen) at 37°C in 5% CO_2_ during image acquisition and microscope acquisition parameters for all images were identical. Lysosome number per cell was blindly quantified and normalized by cell area.

For DQ-BSA assays, DIV7 primary cortical neurons were incubated with 10μg ml^-1^ DQ Green BSA (ThermoFisher) at 37°C, washed with PBS twice, and fixed with 4% PFA for 10 minutes at room temperature. After fixation, cells were washed with PBS three times, protected from light, and cover slips were mounted on glass slides using Vectashield.

Unless otherwise noted, imaging was performed using a Zeiss LSM880 confocal microscope or Zeiss LSM710 confocal microscope and processed with ImageJ software (National Institutes of Health). Number of NeuN-positive layer V cortical neurons was quantified as previously described (34), with minor modifications. Briefly, following acquisition and stitching of a 3×3 tiled array of 20x images, stitched images were de-identified and randomized. The polygon selection tool of Fiji (60) was used to outline layer V, and the area of the selection was measured using the ‘measure’ tool and recorded. Then, the area outside of the selection was removed using the ‘clear selection’ tool and a threshold was applied using the ‘otsu’ algorithm. Overlapping nuclei were separated using the ‘watershed’ tool and nuclei larger than 50μm^2^ with a circularity between 0.2-1.0 were counted using the ‘analyze particles’ tool. Number of nuclei was divided by the area of the polygon selection to determine density of nuclei. For all histological analyses, at least three animals were examined, with at least 3-9 30μm sections from comparable brain/spinal cord regions quantified per animal. For Lamp1 quantification in spinal cords, only cells with soma size greater than 200μm^2^ were counted. The experimenter was blind to animal genotype and ASO treatment during image acquisition and quantification.

The following primary antibodies used were: rat anti-Lamp2 (Abcam; ab25339; 1:400), mouse anti-Cathepsin D (Abcam; ab6313; 1:200), goat anti-ChAT (Millipore; AB144P; 1:500), rabbit anti-Lamp1 (Abcam; ab24170; 1:400), chicken anti-GFAP (Abcam; ab4674; 1:2,000), mouse anti-NeuN (Millipore; MAB377; 1:200), mouse anti-SMI32 (Covance; SMI-32P; 1:1,000), mouse anti-Flag (Sigma-Aldrich; F3165; 1:2000), and chicken anti-GFP (Abcam; ab13970; 1:3,000).

### Luciferase reporter assay

Cells were transfected with 4XCLEAR-Luciferase reporter construct (Addgene #66800)(11) and internal control pRL-TK vector (Promega) plasmids using Lipofectamine 2000. Transfected cells were cultured in DMEM supplemented with 10% FBS for two days. Cells were then collected, lysed, and subjected to dual luciferase reporter assay using Dual-luciferase Reporter Assay System (Promega) and GloMax 2020 luminometer (Promega) according to manufacturer’s instruction. Luciferase activity was normalized by dividing the *firefly* luciferase activity from the 4XCLEAR-Luciferase construct by the *Renilla* luciferase activity from pRL-TK vector. All experiments were repeated in triplicate and performed at least three times.

### Antisense oligonucleotide (ASO) generation and administration

Synthesis and purification of Control ASO (CCTATAGGACTATCCAGGAA) or *Nlk*-targeting ASO (GTCACAGTACAGCCTGGATC) were performed as previously described (61). The MOE-gapmer ASOs are 20 nucleotides in length, wherein the central gap segment comprising ten 2’-deoxyribonucleotides that are flanked on the 5’ and 3’ wings by five 2’MOE modified nucleotides. Internucleotide linkages are phosphorothiorate interspersed with phosphodiester, and all cytosine residues are 5’ methylcytosines. ASOs were delivered intracerebroventricularly (ICV) through the right hemisphere (2mm anterior to lambdoid suture, 1mm lateral from sagittal suture, 2mm deep) of hypothermia-induced anaesthetized mice at P1 using 10ml Model 1701 RN Syringe with a 33-gauge, 0.375 inch, custom point style 4, 45° angle needle (Hamilton).

### Animal behavioral assessment

Grip strength was assessed in P84 *Prp-TDP*^*A315T*^ animals and littermate controls using a Chatillon grip strength meter (Columbus Instruments). Six measurements were taken for forelimb or forelimb with hindlimb grip strength per animal. The maximum and minimum force measurements were discarded and the remaining four were averaged and the average was recorded for each animal. The experimenter was blind to animal genotype and ASO treatment.

Motor behavior of *Thy1-TDP*^*Tg*^ animals and littermate controls was examined at P14, P16, P18, P20, and P22 as previously described (34, 39), with minor modifications. Briefly, animals were assessed for gait impairment, tremor, kyphosis, and hindlimb clasping. To assess gait impairment, tremor, and kyphosis, animals were placed into a new cage on a flat, but textured plastic surface. For gait impairment, a score between 0 and 4 was assigned based on the following criteria: normal movement (score of 0); grossly normal movement with a mild tremor or limp (score of 1); severe tremor, limp, or lowered pelvis during movement (score of 2); difficulty moving forward with frequent falls, but is able to right itself within 30 seconds of falling (score of 3); unable to right itself in three consecutive 30 second trials (score of 4), also considered the humane endpoint for euthanasia. For tremor, a score between 0 and 3 was assigned based on the following criteria: no tremor (score of 0); mild tremor while moving (score of 1); severe jerking tremor while moving (score of 2); severe tremor at rest and during movement (score of 3). For determining kyphosis score, a score between 0 and 3 was assigned based on the following criteria: no observable kyphosis (score of 0); mild kyphosis at rest but able to straighten spine completely during movement (score of 1); mild kyphosis at rest and unable to straighten spine completely during movement (score of 2); severe kyphosis at rest that is maintained during movement (score of 3).

To assess hindlimb clasping, animals were suspended by the base of their tail and the hindlimb positioning was observed for 10 seconds. A score between 0 and 3 was assigned based on the following criteria: hindlimbs extended outwards for 50% or more of trial (score of 0); one hindlimb pointing inward towards the abdomen for 50% or more of trial (score of 1); both hindlimbs partially pointed towards the abdomen for 50% or more of the trial (score of 2); both hindlimbs completely clasped inwards together against the abdomen for 50% or more of the trial (score of 3). Composite motor behavior scores were calculated by summing gait impairment, kyphosis, tremor, and hindlimb clasping scores (max score of 13) for each animal. For animals that reached their humane endpoint prior to P22 measurements, the scores for gait impairment, tremor, and hindlimb clasping at time of endpoint were also reported as scores for the remaining recordings for those animals. For animals sacrificed at P19 for pathology studies, no scores were reported for P20 and P22 timepoints. The experimenter was blind to animal genotype and ASO treatment.

### Statistical analysis

Unless otherwise noted, all data were analyzed by two-tailed, unpaired Student’s t-test (two experimental groups) or one-way ANOVA with multiple comparisons (more than two experimental groups) to determine statistical significance between samples using Prism 7. P<0.05 was considered to be statistically significant. Graphs were plotted using Prism 7 and heatmaps and volcano plots were plotted using R version 3.3.3.

## Data availability

RNA-data FASTQ files and processed normalized output have been deposited into GEO under accession #xxxxx. The data, protocols, and code that were generated and support the findings of this study are available from the corresponding author upon reasonable request. The ASOs used in this study are produced by Ionis Pharmaceuticals, a for-profit company.

## Supplementary Materials

Supplemental Figure 1. Generation of isogenic *Nlk* N2a cell lines using CRISPR/Cas9.

Supplemental Figure 2. RNA-seq of *Nlk* KO cells reveals transcriptional changes associated with the autophagy-lysosome pathway.

Supplemental Figure 3. Nlk inhibits protein clearance through the autophagy-lysosome pathway.

Supplemental Figure 4. Generation of *NLK*-deficient human iPSC-derived motor neurons.

Supplemental Figure 5. Nlk does not physically interact with TDP-43.

Supplemental Figure 6. Reduction of *Nlk* improves survival and delays onset of disease in *Prp- TDP*^*A315T/+*^ mice.

Supplemental Figure 7. Pharmacological reduction of *Nlk* does not induce gliosis *in vivo*.

Supplemental Figure 8. ASOs targeting *Nlk* ameliorate motor behavior deficits in *Thy1-TDP*^*Tg/Tg*^ mice.

Supplemental Figure 9. ASOs targeting *Nlk* ameliorate motor behavior deficits in high fat/gel diet-fed *Thy1-TDP*^*Tg/Tg*^ mice.

Supplemental Figure 10. ASOs targeting *Nlk* rescue pathology deficits in two TDP-43 mouse models.

Supplemental Table 1. Differential gene expression from RNA-seq of *Nlk* KO N2a cells.

Supplemental Video 1. ASOs targeting *Nlk* partially rescue motor impairment in *Thy1-TDP*^*Tg/Tg*^ mice.

## Acknowledgments

We thank Dr. Terri Driessen for advising on RNA-seq analysis, all members of the Lim laboratory and Dr. Shawn Ferguson for useful feedback, critiques, and comments, and the Yale Center for Genome Analysis (YCGA) for performing RNA sequencing.

## Author contributions

L.T., H.K., and J.L. conceived and designed this study. H.K. generated *Nlk* KO N2a cells and performed *in vitro* TDP-43, autophagy, and primary cortical neuron experiments. J.P. performed N2a LysoTracker microscopy and L.T. and J.Y. quantified images. P.J.L. performed bioinformatic analysis of RNA-seq data and generated heatmaps and volcano plots. L.T. and J.L. performed mouse survival studies. L.T. and Y.X. generated mutant *NLK* hiPSC lines and performed human iPSC-derived motor neuron experiments. L.T., with help from H.J., performed *Nlk* genetic reduction mouse experiments. A.S., F.R., and P.J. generated ASOs and advised on experimental design for ASO experiments. L.T. performed mouse ASO experiments, with help from P.J.L., K.L., J.Y., H.R., C.L., and S.M.T for mouse behavior, dissections, immunostaining, protein, and RNA extraction. P.J.L. maintained the mouse colony and genotyped animals. L.T. and J.L. wrote the manuscript.

## Author Information

The authors declare competing financial interests: A.S., F.R., and P.J. are employed by Ionis Pharmaceuticals, a for-profit company that develops ASO technologies. The other authors declare no competing financial interest.

## Data and materials availability

RNA-data FASTQ files and processed normalized output have been deposited into GEO under accession #xxxxx. The data, protocols, and code that were generated and support the findings of this study are available from the corresponding author upon reasonable request. The ASOs used in this study are produced by Ionis Pharmaceuticals, a for-profit company. Correspondence and requests for materials should be addressed to.

## Supplementary Materials

**Supplemental Figure 1.**
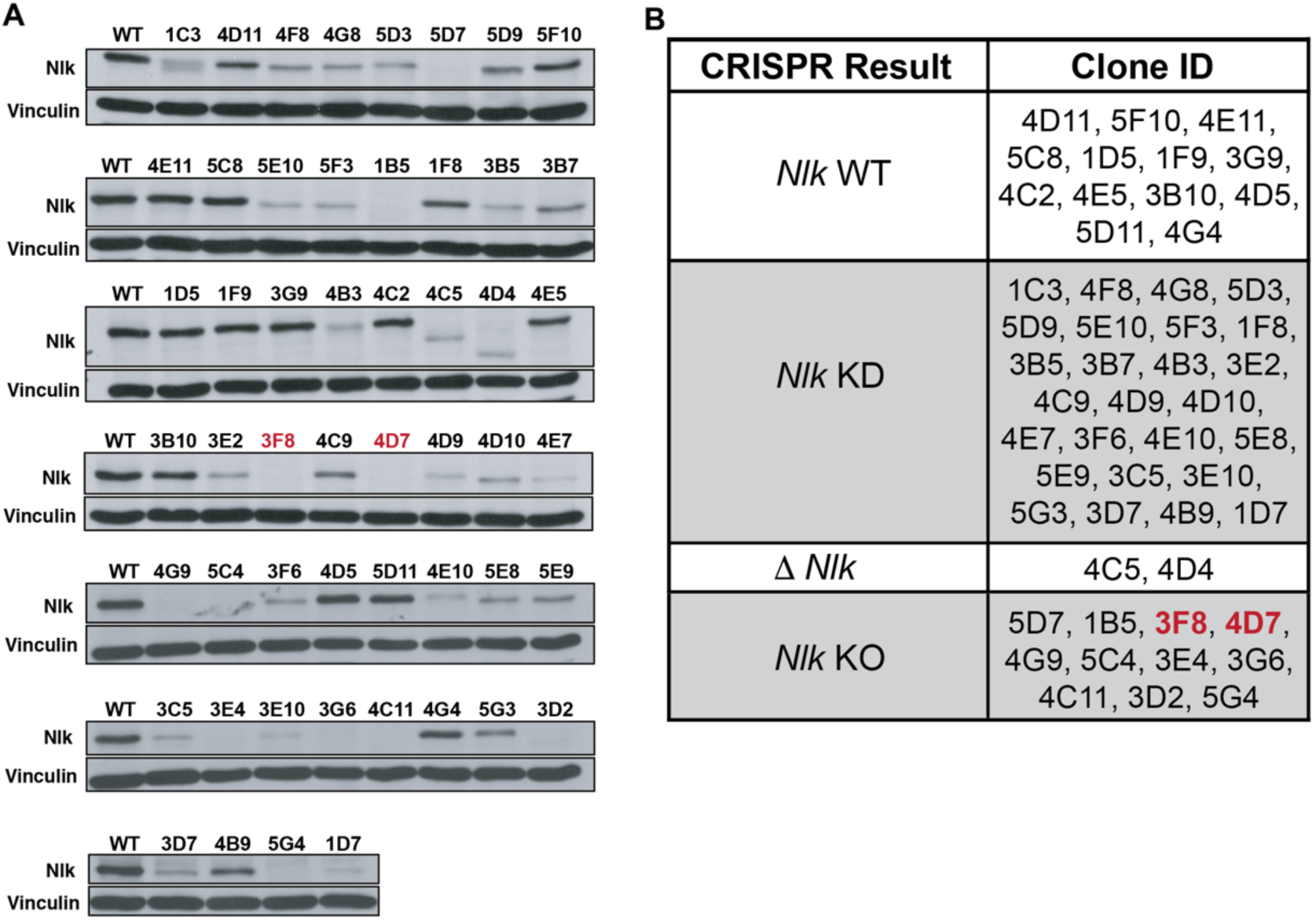
Generation of isogenic *Nlk* N2a cell lines using CRISPR/Cas9. (**A**) Western blots of protein lysates detecting the degree of mutagenesis in individual clones isolated following dual guide RNA targeting of Cas9 nickase to *Nlk*. (**B**) Summary of genotypes of *Nlk* CRISPR clones determined remaining levels of Nlk protein. WT= wild-type, KD= knock-down, KO= knock-out. Δ= partial deletion. *Nlk* KO clones 3F8 and 4D7 were selected for further experiments.

**Supplemental Figure 2.**
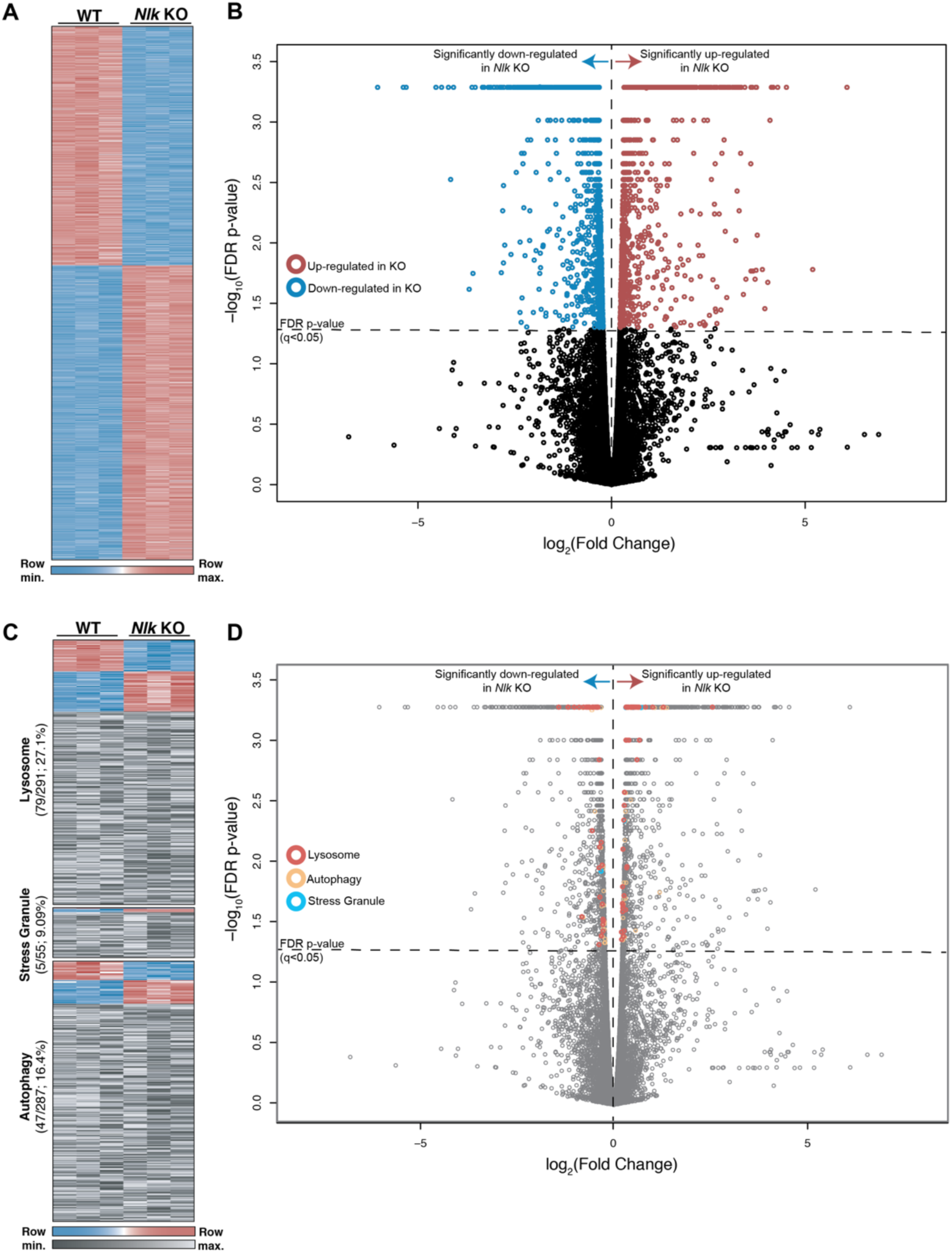
RNA-seq of *Nlk* KO cells reveals transcriptional changes associated with the autophagy-lysosome pathway. (**A**) Heatmap of all significantly altered genes from RNA-seq analysis of *Nlk* KO N2a cells. (**B**) Volcano plot of RNA-seq results displaying all significantly up-regulated (red) and down-regulated (blue) genes. (**C**) Individual heatmaps displaying all significantly altered (blue/red) and not significantly altered (gray) annotated lysosome, autophagy, stress granule-associated genes with detectable expression. (**D**) Volcano plot of RNA-seq results with differentially-expressed lysosome, autophagy, and stress granule-associated genes.

**Supplemental Figure 3.**
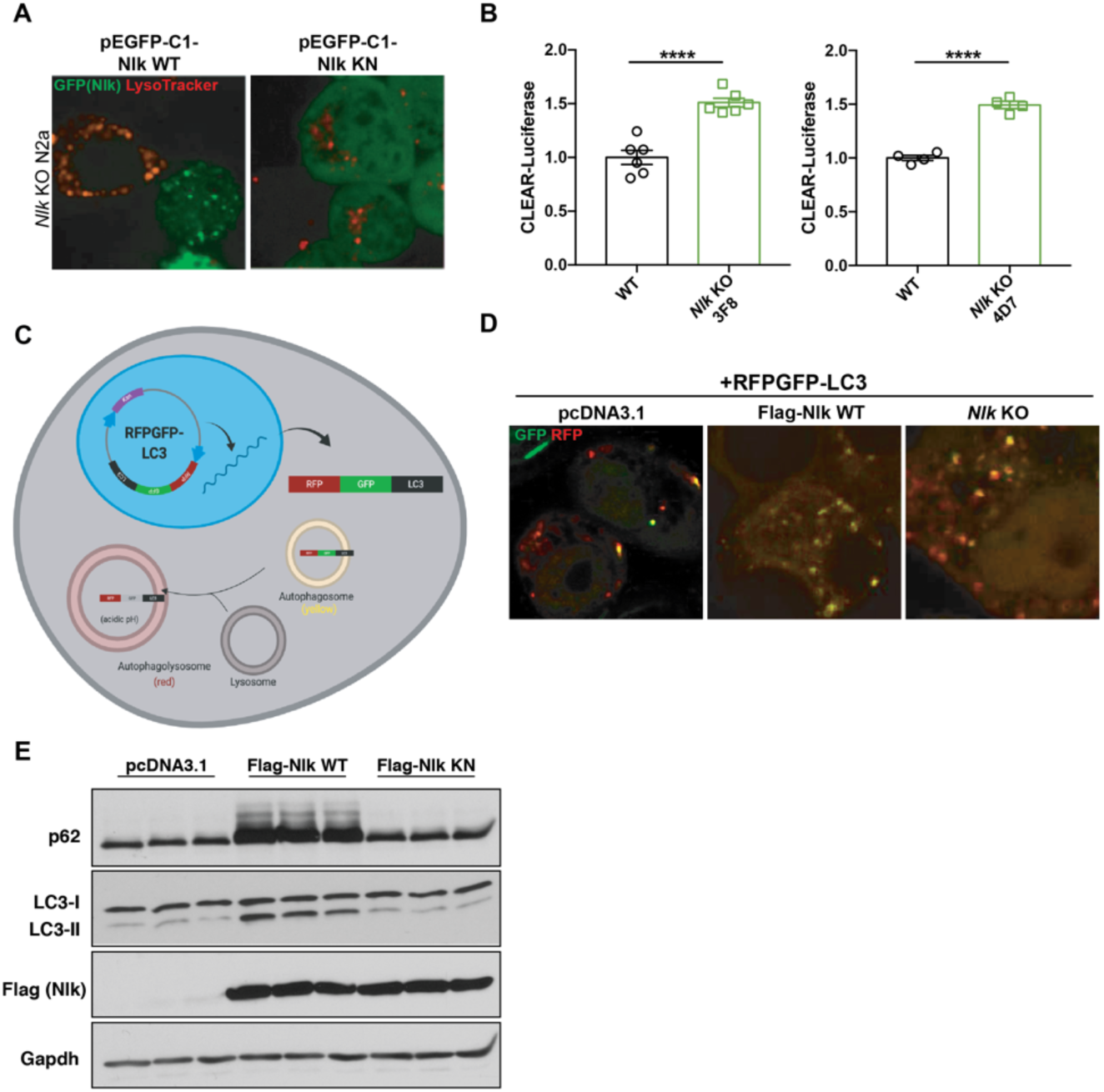
Nlk inhibits protein clearance through the autophagy-lysosome pathway. (**A**) LysoTracker imaging of *Nlk* KO N2a cells transfected with WT or KN Nlk. Cells successfully transfected with WT Nlk (green) contained fewer LysoTracker-positive vesicles compared to non-transfected cells or cells transfected with kinase-inactive Nlk (Nlk KN). (**B**) CLEAR-Luciferase assay in two independent *Nlk* KO N2a clones (pooled in Figure 1K). Two tailed t-tests were performed and mean±s.e.m are displayed. ***p<0.001. (**C**) Schematic detailing the tandem fluorescent-tagged RFPGFP-LC3. Transfected cells express the RFPGFP-LC3 fusion protein, which is trafficked into autophagosomes, where red and green co-localized fluorescence will be observed. Upon fusion with a lysosome, the green fluorescence is quenched by the pH of the lysosome and red-labeled vesicles can be detected. (**D**) Overexpression of WT Nlk reduced the number of autophagolysosomes (red), while *Nlk* KO cells have increased number. (**E**) Western blots showed accumulation of p62 and LC3-II due to the failure of lysosome-dependent degradation in cells with WT Nlk overexpression. Two-tailed t-tests were performed between the two genotypes and mean±s.e.m are displayed. ****p<0.0001.

**Supplemental Figure 4.**
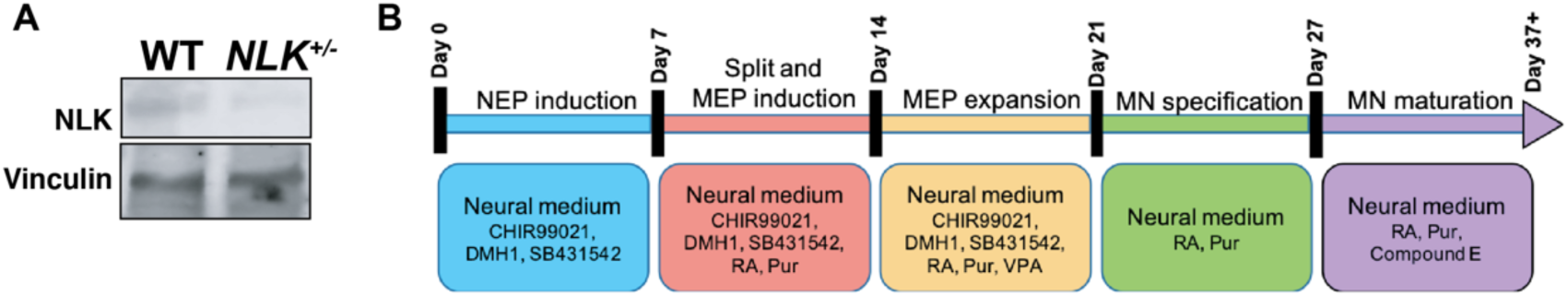
Generation of *NLK*-deficient human iPSC-derived motor neurons. (**A**) Western blots of day 38 WT and isogenic *Nlk* heterozygous clones differentiated into motor neurons. (**B**) Schematic detailing the protocol for generation of spinal motor neurons from iPSCs.

**Supplemental Figure 5.**
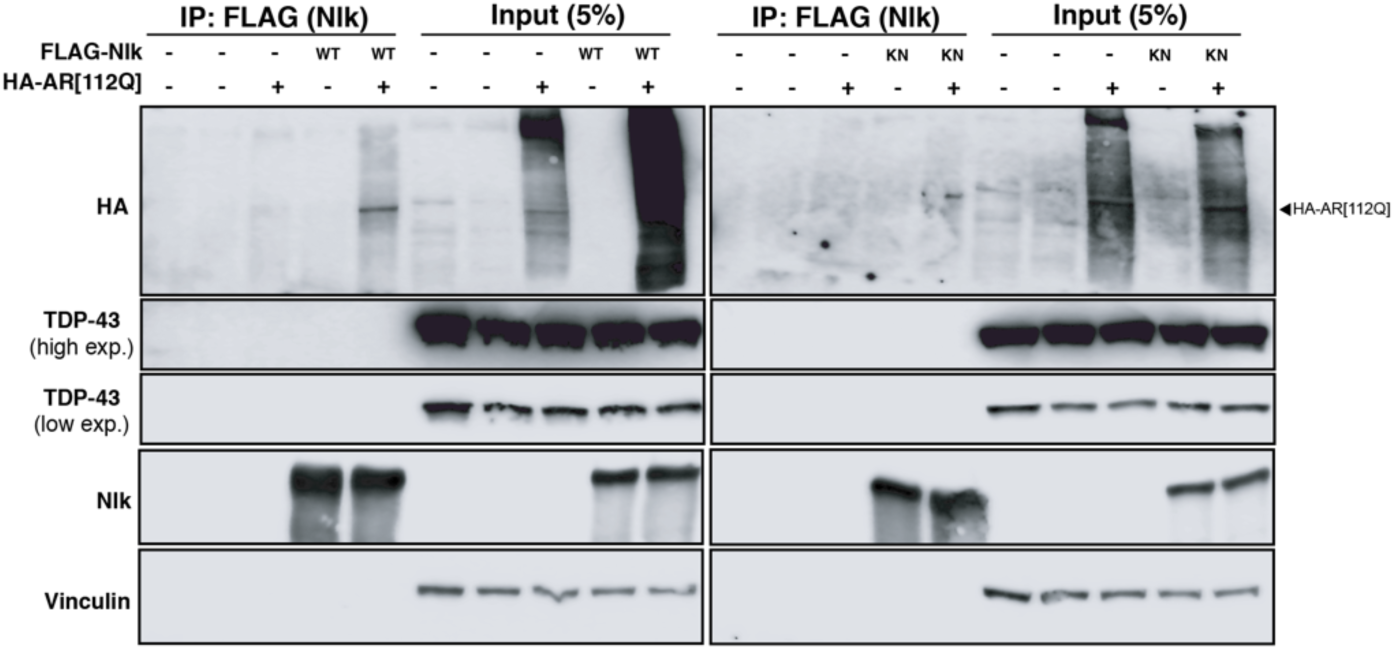
Nlk does not physically interact with TDP-43. Co-immunoprecipitation showing that Nlk-WT and Nlk-KN interact with AR but not with TDP-43. Interaction of Nlk with AR serves as a positive-control (see ref. 10).

**Supplemental Figure 6.**
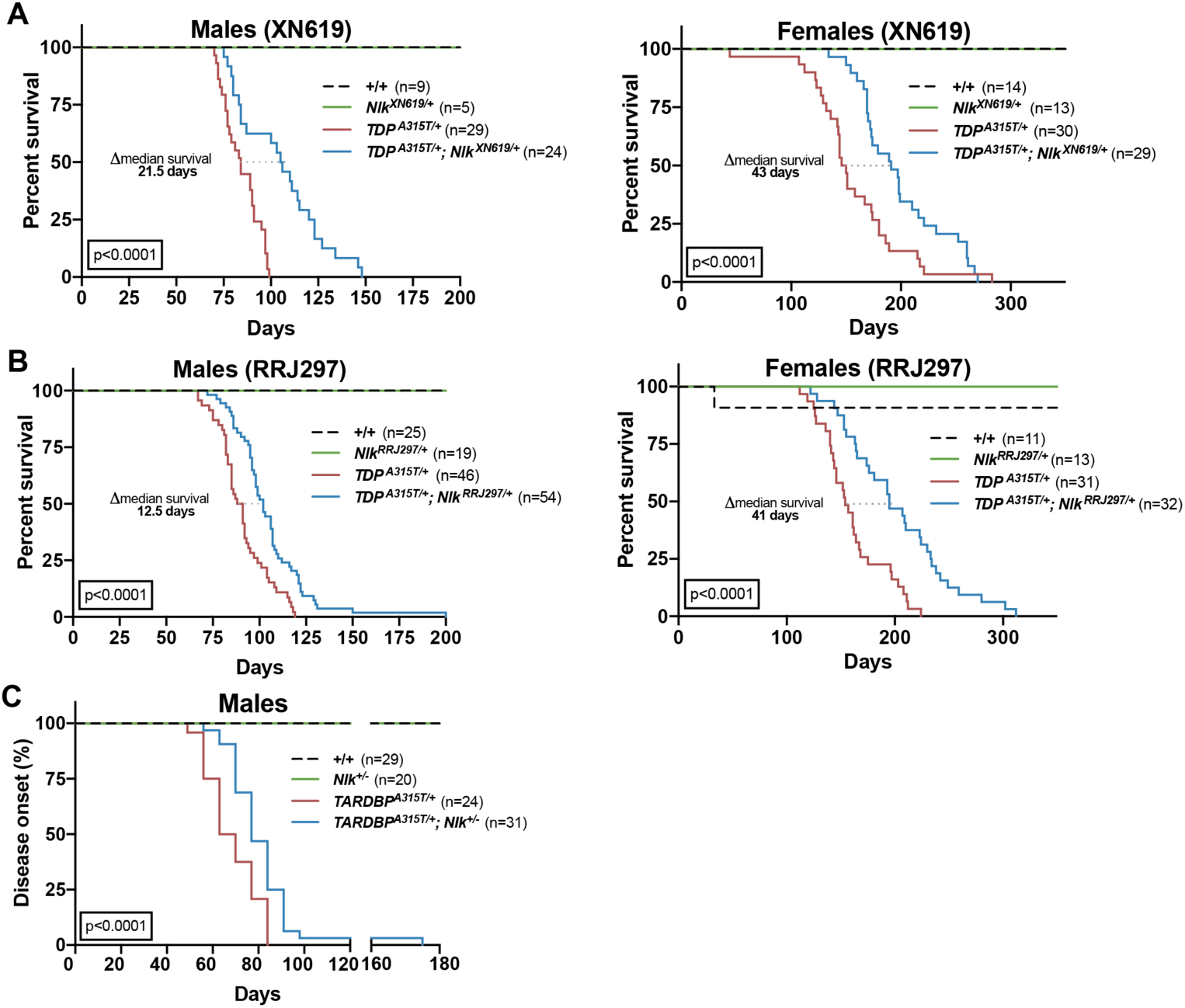
Reduction of *Nlk* improves survival and delays onset of disease in *Prp- TDP*^*A315T/+*^ mice. (**A**) Kaplan-Meier survival curves showing 50% reduction of *Nlk* by crossing *Prp-TDP*^*A315T/+*^ mice with *Nlk*^*XN619/+*^ improved male and female animal survival. Curves were compared by long-rank test. (**B**) Kaplan-Meier survival curves showing 50% reduction of *Nlk* by crossing *Prp-TDP*^*A315T/+*^ mice with *Nlk*^*RRJ297/+*^ improved male and female animal survival. Curves were compared by log-rank test. (**C**) Kaplan-Meier curves showing 50% reduction of *Nlk* by crossing *Prp-TDP*^*A315T/+*^ mice with *Nlk*^*+/-*^ significantly slowed time to disease onset (defined by the earliest time point in which weight gain was no longer observed) in male mice.

**Supplemental Figure 7.**
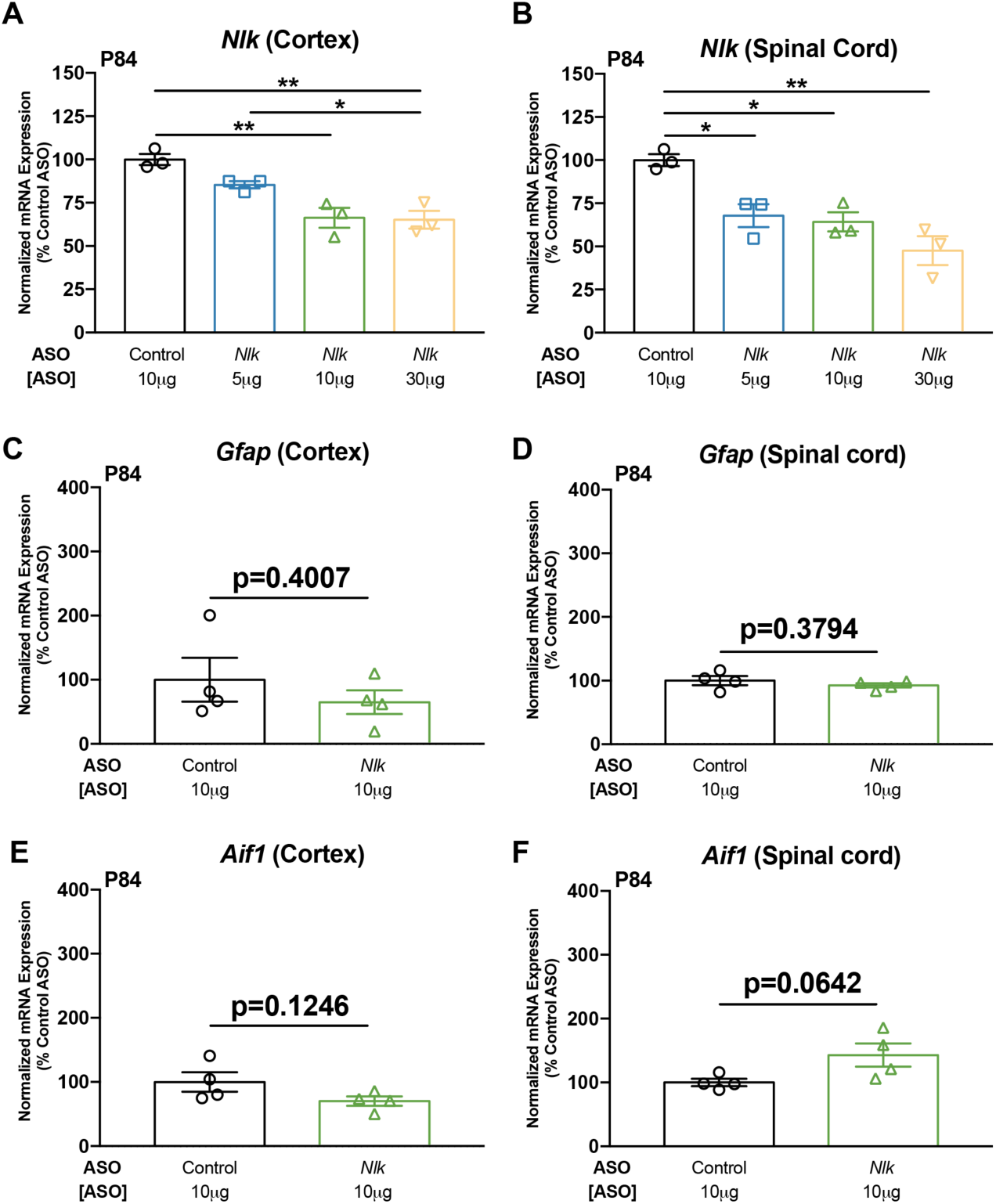
Reduction of *Nlk* does not induce gliosis *in vivo*. (**A-F**) P1 delivery of *Nlk* ASOs reduced *Nlk* mRNA levels in a dose-dependent manner (**A**,**B**) without inducing an overt astrogliosis (**C**,**D**) or microgliosis (**E**,**F**) in the cortex and spinal cord of P84 animals. One-way ANOVA analyses were performed to compare effects of varying ASO dosage. Two-tailed t-tests were performed to compare conditions unless otherwise noted and mean±s.e.m are displayed. *p<0.05, **p<0.01.

**Supplemental Figure 8.**
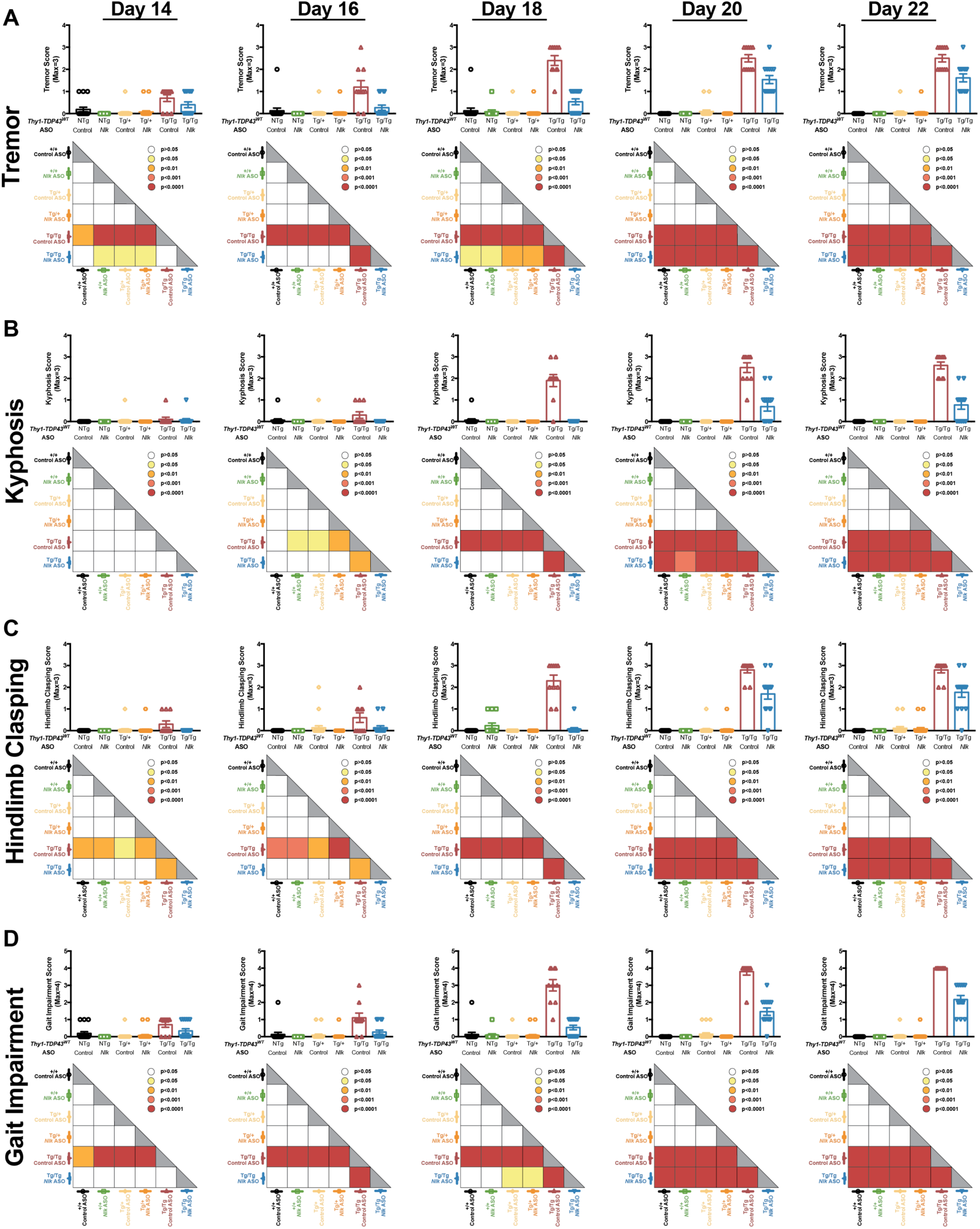
ASOs targeting *Nlk* ameliorate motor behavior deficits in *Thy1-TDP*^*Tg/Tg*^ mice. (**A-D**), Bar plots showing tremor (**A**), kyphosis (**B**), hindlimb clasping (**C**), and gait impairment (**D**) scores for ASO-injected animals on days 14, 16, 18, 20, and 22. Higher score numbers indicated increasing severity (see Methods). Results of statistical analyses are reported in each triangle below the corresponding bar graph. One-way ANOVA analyses were performed to compare each genotype/condition on each individual day and reported in the statistics triangle below each bar plot.

**Supplemental Figure 9.**
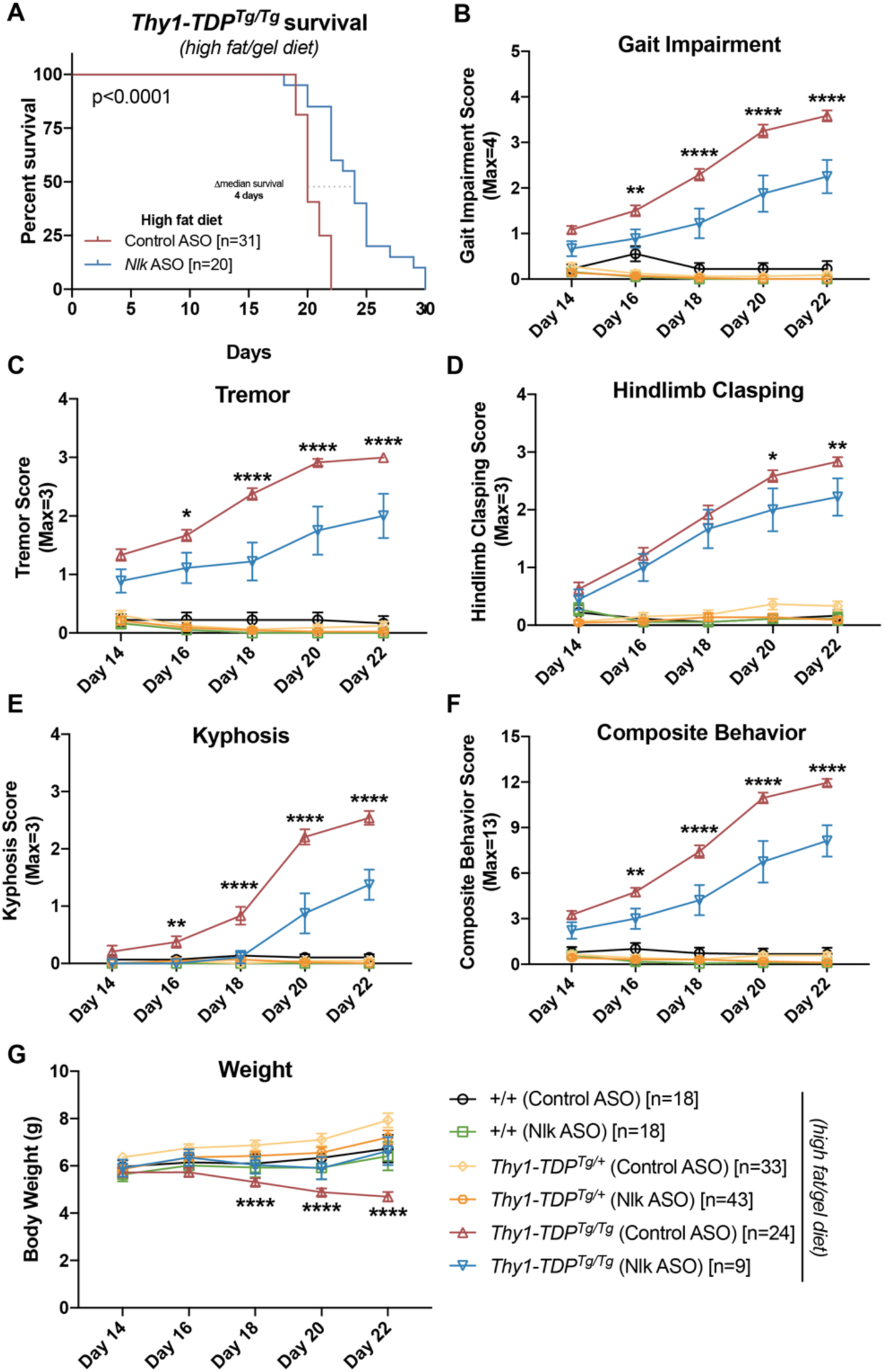
ASOs targeting *Nlk* ameliorate motor behavior deficits in high fat/gel diet-fed *Thy1-TDP*^*Tg/Tg*^ mice. (**A**) Kaplan-Meier survival curves showing administration of 10μg *Nlk* ASO at P1 increased *Thy1-TDP*^*Tg/Tg*^ animal survival. Curves were compared by log-rank test. (**B-G**) *Nlk* ASO administration reduced gait impairment (**B**), tremor (**C**), hindlimb clasping (**D**), kyphosis (**E**), composite motor score (**F**), and prevented weight loss (**G**) in *Thy1-TDP*^*Tg/Tg*^ animals between P14-P22. One-way ANOVA analyses were performed to compare all listed genotypes/treatments per day unless otherwise noted and mean±s.e.m are displayed. *p<0.05, **p<0.01, ****p<0.0001.

**Supplemental Figure 10.**
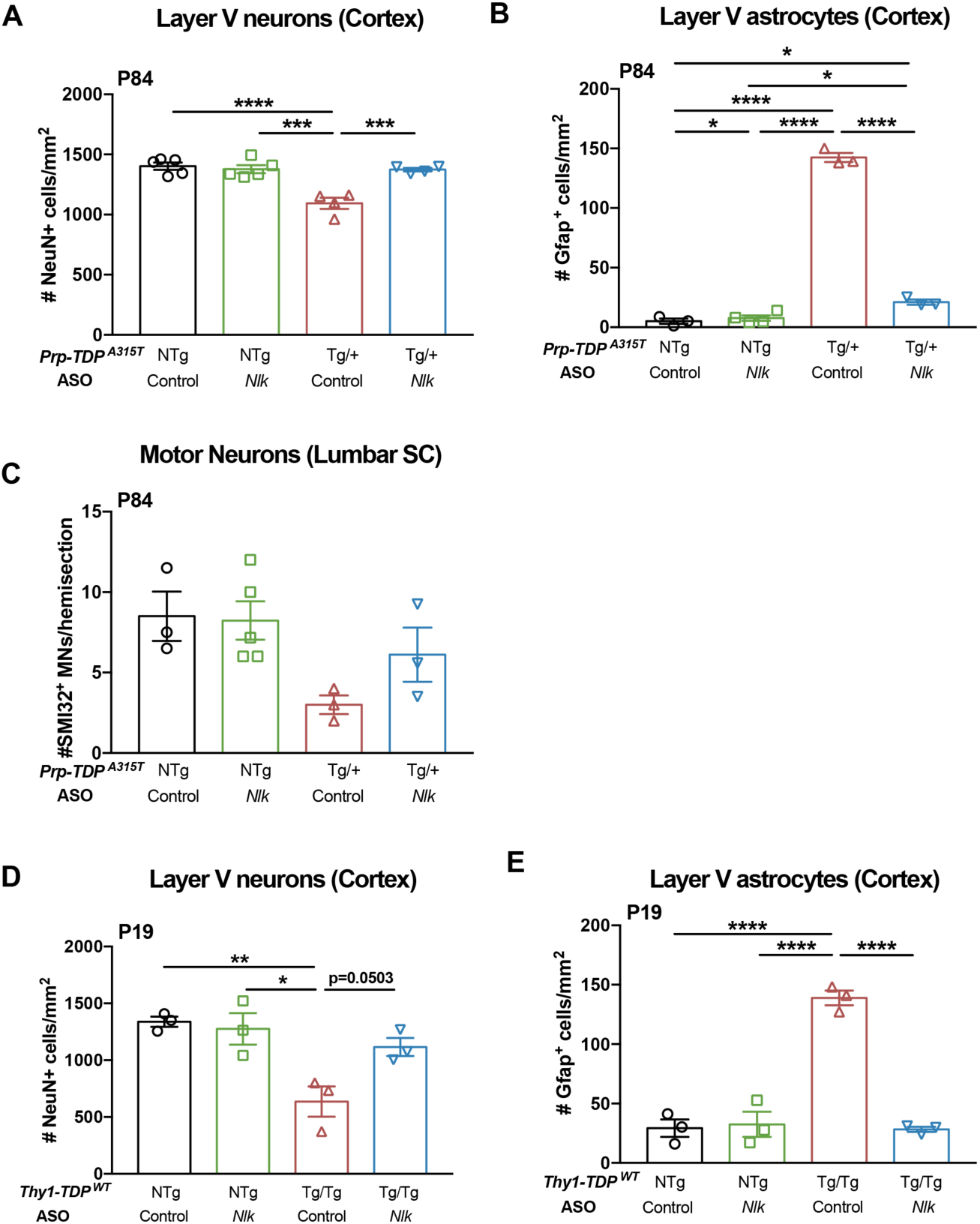
ASOs targeting *Nlk* rescue pathology deficits in two TDP-43 mouse models. (**A-C**) Administration of 10μg of *Nlk* ASO at P1 rescued loss of layer V cortical neurons (**A**; n=animals), layer V astrogliosis (**B**; n=animals), and lumbar spinal cord (SC) motor neuron loss (**C**; n=animals) in P84 *Prp-TDP*^*A315T/+*^ male mice. (**D**,**E**) 10μg of *Nlk* ASO at P1 rescued loss of layer V cortical neurons (**D**; n=animals) and layer V astrogliosis (**E**; n=animals) in *Thy1-TDP*^*Tg/Tg*^ mice at P19. One-way ANOVA analyses were performed to compare all listed genotypes/conditions and mean±s.e.m are displayed. *p<0.05, **p<0.01, ***p<0.001, ****p<0.0001.

**Supplemental Table 1. Differential gene expression from RNA-seq of *Nlk* KO N2a cells**. Processed and normalized differential gene expression data from RNA-seq comparing transcriptomes of *Nlk* KO N2a cells to isogenic controls. Mean expression data from triplicates were compared and FDR-adjusted p-value (q<0.05) was used to determine significance.

**Supplemental Video 1. ASOs targeting *Nlk* partially rescue motor impairment in *Thy1-TDP***^***Tg/Tg***^ **mice**. *Thy1-TDP*^*Tg/Tg*^ mice injected with control ASO displayed a rapid stereotyped decline between P18 and P19, necessitating euthanasia. *Thy1-TDP*^*Tg/Tg*^ mice injected with 10μg of *Nlk* ASO at P1 retained motor abilities beyond P19.

